# Hippocampal-cortical interactions in the consolidation of social memory

**DOI:** 10.1101/2024.10.10.617719

**Authors:** Gaeun Park, Min Seok Kim, Young-Beom Lee, Soonho Shin, Doyun Lee, Sang Jeong Kim, Yong-Seok Lee

## Abstract

Episodic memories are initially encoded in the hippocampus and subsequently undergo systems consolidation into the neocortex. The nature of memory stored in the hippocampus and neocortex also differs, with the cortex shown to encode memories in more generalized forms. Although several brain regions are known to encode social information, the specific cortical regions and circuits involved in the consolidation of social memory and the nature of the information encoded in the cortex remain unclear. Using *in vivo* Ca^2+^ imaging and optogenetic manipulations, we found that infralimbic (IL) neurons projecting to the nucleus accumbens shell (IL^→NAcSh^) neurons store consolidated social memory. Inactivating IL^→NAcSh^ neurons that responded to a familiar conspecific impaired the recognition of other familiar mice including littermates, demonstrating that these neuronal activities support social familiarity. Furthermore, inactivating hippocampal ventral CA1 neurons projecting to the IL disrupted the consolidation of memory for newly familiarized mice while sparing the recognition of littermates. These findings demonstrate the critical role of hippocampal-cortical interactions in the consolidation of social memory.

## Introduction

Episodic memory is initially encoded and stored in the hippocampus and then consolidated into the cortex through intricate hippocampal-cortical interactions^1,2^. Prevalent models on the organization of memory such as the complementary learning systems theory propose that the interaction between the hippocampus and cortex during offline period is crucial for memory storage, especially for recognition memory^3^. In addition, a number of memory models have proposed that different brain regions encode distinct aspects of memory^4,5^. For example, the cortex stores memory in more generalized or abstract forms compared to the hippocampus^6^.

Social memory is the ability to recognize and remember conspecific, consists of the detection of familiarity and the recollection of social episode^7,8^. The hippocampal dorsal CA2 (dCA2) is considered as a hub region for social memory in rodents, with a recent study showing that the tuned geometries of dCA2 representations can support both familiarity and recollection ^8–11^. The neural projections from the lateral entorhinal cortex were shown to convey social information to the dCA2, which in turn send its projections to the ventral CA1 (vCA1)^11,12^. vCA1 to the nucleus accumbens shell (NAcSh) projections have been also shown to be critical for social memory^13,14^. While vCA1 to mPFC projections have been implicated in social memory^15,16^, the precise role of hippocampal-cortical interactions in social memory consolidation remains largely unknown. Moreover, whether the mPFC stores social memory in generalized forms is unclear.

Neural representations of social information have been reported from the PFC neurons, encoding social *vs.* nonsocial information which include relevant information such as sex or hierarchy^17–20^. Accordingly, social representations in the PFC are impaired in mouse models of autism spectrum disorder^20,21^. It has also been shown that learning promotes task-relevant tuning and alterations in the dimensionality of neural representations in the PFC^22–24^. However, our understanding on social representations in the PFC is still limited. Previously, we reported that a subpopulation of mPFC infralimbic (IL) neurons projecting to the NAcSh are activated by familiar mice and chemogenetic suppression of the activity of these neurons impairs social recognition, suggesting that the IL^→NAcSh^ neurons is critical for processing social memory^25^. However, at which stage of memory processing that these neurons are engaged as well as the nature of social information encoded in these neurons remain unclear.

In this study, we aimed to test our hypothesis that IL^→NAcSh^ neurons stores consolidated social memory in a generalized form via hippocampal-cortical interactions. To investigate this, we focused on the role of vCA1-IL-NAcSh circuit in social memory consolidation in male mice by using a combination of *in vivo* Ca^2+^ imaging, viral tracing and tagging, as well as optogenetic and chemogenetic manipulations.

## Results

### Inactivation of IL^→NAcSh^ neurons impairs social memory retrieval

Previously, we have shown that IL^→NAcSh^ neurons are critically involved in social recognition^25^. However, the specific temporal window during which IL^→NAcSh^ neurons are engaged remains unclear. The social familiarization/recognition task was used to investigate social recognition memory in male mice (Fig. 1a). During social familiarization, each subject mouse was exposed to a novel conspecific (F_N_) which eventually became a familiar mouse (F). IL^→NAcSh^ neurons were inactivated in different phrases of the social familiarization/recognition task by using optogenetic or chemogenetic manipulations (Fig. 1b). To inactivate IL^→NAcSh^ neurons during either the familiarization (encoding) or recognition (retrieval) phase, halorhodopsin (NpHR) or yellow fluorescent protein (YFP) was expressed in the IL while optic fibers were bilaterally targeted in the NAcSh (Fig. 1c). Optogenetic inactivation of the IL^→NAcSh^ neurons during social familiarization on day 1 did not affect either social familiarization or recognition in both NpHR and YFP-expressing mice (Fig. 1d and Supplementary Fig. 1a). On day 2, both groups showed significantly longer interaction times with the novel conspecifics (N) than either the familiar conspecific (F) or the littermate (L). When these neurons were chemogenetically inactivated during the offline phase after the social familiarization, social preference for novel conspecifics was also intact during the social recognition session on day 2 (Figs. 1e and 1f, Supplementary Fig. 1b). However, when we inactivated these neurons during the social recognition on day 2, NpHR-expressing mice exhibited impaired social recognition, while YFP-expressing mice showed a preference for the novel conspecific (Fig. 1g). The same mice showed normal social recognition without optogenetic manipulations (Supplementary Fig. 1c). These results indicate that IL^→NAcSh^ neuronal activity is required for the retrieval of social memory. In addition, optogenetic manipulation did not affect locomotive activities or real-time place preference (Supplementary Figs. 1d and 1e).

**Figure 1.**
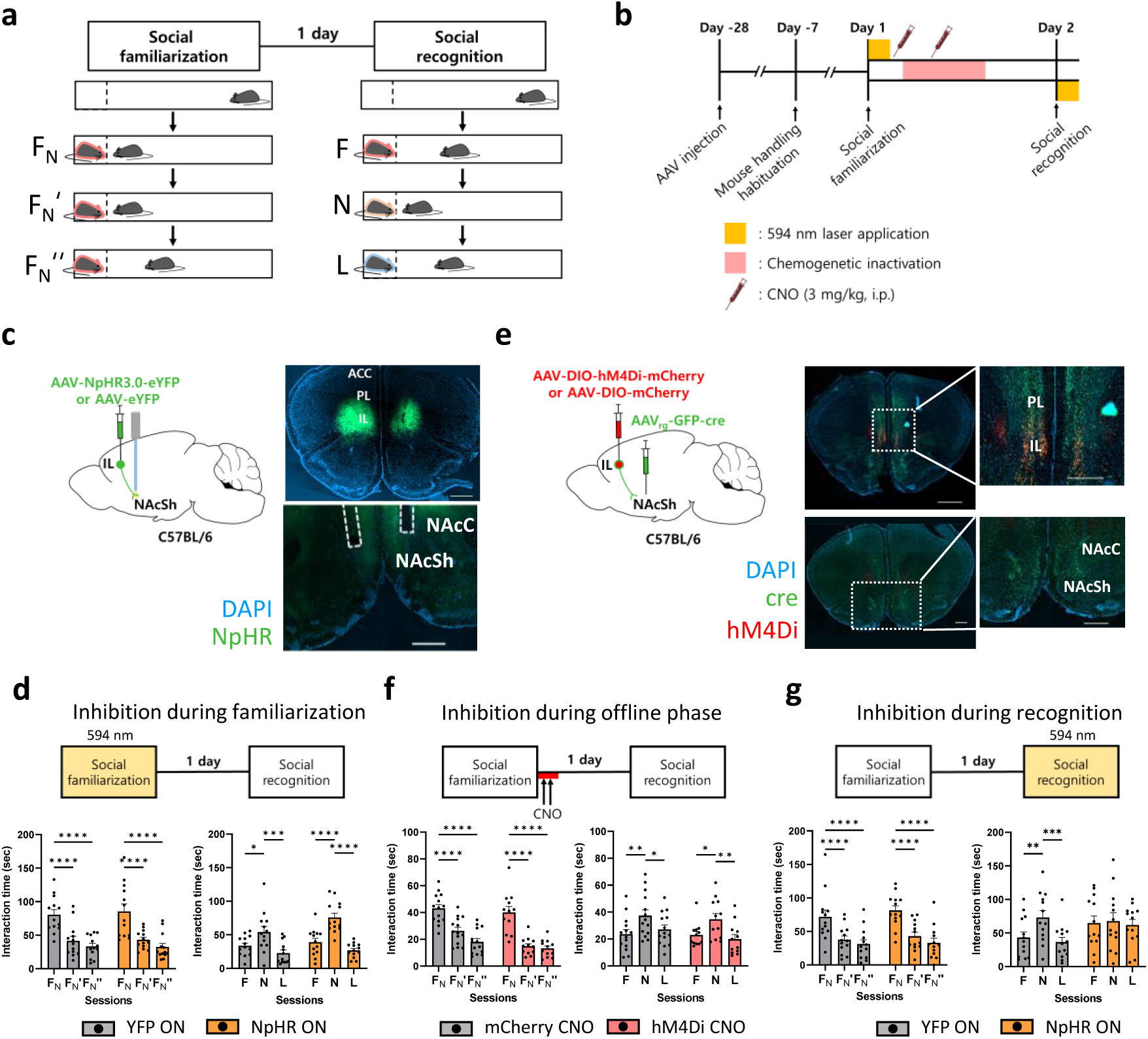
Inactivation of IL^→NAcSh^ neurons impairs social memory retrieval. (a) Behavior scheme of the social familiarization/recognition task. During the social familiarization session, the subject mouse interacts with a novel target conspecific (F_N_) 3 times for 5 minutes, with 5 min intervals after 5 minutes of habituation session. The target conspecific was re-defined as F_N_’ and F_N_’’ following repeated interactions. On the next day (social recognition session), the subject mouse interacts with the familiarized social target (F), a novel target (N) and a littermate (L) for 5 min each. (b) Timeline for optogenetic and chemogenetic manipulations of IL^→NAcSh^ neurons. (c) Left: schematic diagram of optogenetically inactivating IL^→NAcSh^ neurons. NpHR3.0 or eYFP was expressed in IL neurons while two 200μm optic fibers were bilaterally implanted in NAcSh. Right: representative image of the mPFC (top, Scale bar: 1000 μm) and NAcSh (bottom, Scale bar: 500 μm) expressing NpHR in neuronal soma and terminal, respectively. (d) Left: both YFP and NpHR mice showed significantly decreased interaction times during the social familiarization session. Two-way ANOVA with repeated measures: YFP, n = 13 mice; NpHR, n = 13 mice; interaction between session and laser effect, F_2,48_ = 0.1333, p = 0.8755. Tukey’s multiple comparison test: YFP, F_N_ versus F_N_’ q_48_ = 6.958, ****p < 0.0001; YFP, F_N_ versus F_N_’’ q_48_ = 8.444, ****p < 0.0001; YFP, F_N_’ versus F_N_’’ q_48_ = 1.486, p = 0.5488; NpHR, F_N_ versus F_N_’ q_48_ = 7.542, ****p <0.0001; NpHR, F_N_ versus F_N_’’ q_48_ = 9.474, ****p < 0.0001; NpHR, F_N_’ versus F_N_’’ q_48_ = 1.931, p = 0.3667. Right: both YFP and NpHR mice showed significantly longer interaction times with novel conspecifics than familiar conspecifics or littermates. Two-way ANOVA with repeated measures: YFP, n = 13 mice; NpHR, n = 13 mice; interaction between target effect and familiarization-laser effect, F_2,48_ = 3.509, *p = 0.0378. Tukey’s multiple comparison test: YFP, F versus N q_12_ = 4.071, ∗p = 0.0344; YFP, F versus L q_12_ = 3.893, *p = 0.0430; YFP, N versus L q_12_ = 8.731, ***p = 0.0001; NpHR, F versus N q_12_ = 12.11, ****p < 0.0001; NpHR, F versus L q_12_ = 3.814, *p = 0.0475; NpHR, N versus L q_12_ = 11.90, ****p < 0.0001. (e) Left: schematic diagram of expressing inhibitory G-protein (Gi)-coupled hM4Di receptor or mCherry selectively in IL^→NAcSh^ neurons by injecting a Cre-dependent hM4Di or mCherry virus in the IL and retrograde eGFP-cre virus in the NAcSh of mice. Right: representative images (Scale bar: 1000 μm) and its magnified images (Scale bar: 500 μm) of IL (top) and NAcSh (bottom). (f) Left: both mCherry and hM4Di mice showed significantly decreased interaction times during the social familiarization. Two-way ANOVA with repeated measures: mCherry, n = 14 mice; hM4Di, n = 12 mice; interaction between session and offline-CNO effect, F_2, 48_ = 2.593, p = 0.0852. Tukey’s multiple comparison test: mCherry, F_N_ versus F_N_’ q_48_ = 8.947, ****p < 0.0001; mCherry, F_N_ versus F_N_’’ q_48_ = 13.35, ****p < 0.0001; mCherry, F_N_’ versus F_N_’’ q_48_ = 4.401, **p = 0.0086; hM4Di, F_N_ versus F_N_’ q_48_ = 12.46, ****p < 0.0001; hM4Di, F_N_ versus F_N_’’ q_48_ = 13.28, ****p <0.0001; hM4Di, F_N_’ versus F_N_’’ q_48_ = 0.8207, p = 0.8313. Right: both mCherry and hM4Di mice showed significantly longer interaction times with novel conspecifics than familiar conspecifics or littermates during the social recognition session. Two-way ANOVA with repeated measures: mCherry, n = 14 mice; hM4Di, n = 12 mice; interaction between session and CNO effect, F_2, 48_ = 0.6990, p = 0.5021. Tukey’s multiple comparison test: mCherry, F versus N q_48_ = 5.015, **p = 0.0025; mCherry, F versus L q_48_ = 1.270, p = 0.6441; mCherry, N versus L q_48_ = 3.744, *p = 0.0289; hM4Di, F versus N q_48_ = 3.938, *p = 0.0205; hM4Di, F versus L q_48_ = 1.053, p = 0.7383; hM4Di, N versus L q_48_ = 4.991, **p = 0.0026. (g) Left: both YFP and NpHR mice showed significantly decreased interaction times during the social familiarization session. Two-way ANOVA with repeated measures: YFP, n = 13 mice; NpHR, n = 13 mice; interaction between session and recognition-laser effect, F_2,48_ = 0.3249, p = 0.7242. Tukey’s multiple comparison test: YFP, F_N_ versus F_N_’ q_48_ = 7.012, ****p < 0.0001; YFP, F_N_ versus F_N_’’ q_48_ = 8.298, ****p < 0.0001; YFP, F_N_’ versus F_N_’’ q_48_ = 1.285, p = 0.6374; NpHR, F_N_ versus F_N_’ q_48_ = 7.901, ****p <0.0001; NpHR, F_N_ versus F_N_’’ q_48_ = 9.907, ****p < 0.0001; NpHR, F_N_’ versus F_N_’’ q_48_ = 2.006, p = 0.3395. Right: NpHR mice showed comparable interaction times with different conspecifics while YFP mice showed a significantly longer interaction times with novel conspecifics, than familiar conspecifics or littermates. Two-way ANOVA with repeated measures: YFP, n = 13 mice; NpHR, n = 13 mice; interaction between target effect and recognition-laser effect, F_2,48_ = 3.752, *p = 0.0306. Tukey’s multiple comparison test: YFP, F versus N q_48_ = 4.896, **p = 0.0032; YFP, F versus L q_48_ = 1.103, p = 0.7172; YFP, N versus L q_48_ = 5.999, ***p = 0.0003; NpHR, F versus N q_48_ = 0.4973, p = 0.9342; NpHR, F versus L q_48_ = 0.4741, p = 0.9400; NpHR, N versus L q_48_ = 0.9714, p = 0.7722.

### IL^→NAcSh^ neurons are activated by familiar conspecifics

Next, we aimed to investigate which aspects of social information are encoded by the IL^→NAcSh^ neurons. Since the inactivation of IL^→NAcSh^ neurons impaired social recognition, we wanted to examine whether these neurons encode the identity or social familiarity of social targets. To investigate social representations in IL^→NAcSh^ neurons, Ca^2+^ activity of this prefrontal subpopulation was monitored by using an open-source one photon miniaturized endoscopic microscope during the social familiarization/recognition task (Supplementary Fig. 2a). A Cre recombinase-dependent genetically encoded Ca^2+^ indicator (GCaMP6f)-expressing Ai148 mice were used to monitor calcium activity in IL^→NAcSh^ neurons, while gradient-index (GRIN) lens attached to a right-angled optical prism was implanted lateral to the prefrontal cortex to image the IL (Fig. 2a). Fluorescence signals (ΔF/F) were obtained and processed to analyze the dynamics of the neuronal activity (Fig. 2b). During the social familiarization, subject mice showed significantly decreased interaction times along consecutive interactions with a novel conspecific (F_N_) (Fig. 2c). During the recognition session on day 2, subject mice showed significantly longer time interacting with the novel conspecific (N) compared to the familiar conspecific (F) or the littermate (L). They also exhibited longer interaction bouts with the novel mouse than with the familiar or littermate mice (Fig. 2c and Supplementary Fig. 2b). This suggests that the miniscope imaging did not affect performances in the social familiarization/recognition task. We longitudinally imaged Ca^2+^ signals during the task and over 80% of GCaMP6f-expressing neurons were successfully registered across days (Supplementary Figs. 2c-2e). Average frequencies of Ca^2+^ transients of the habituation sessions were comparable between the social familiarization and recognition (Supplementary Fig. 2f).

**Figure 2.**
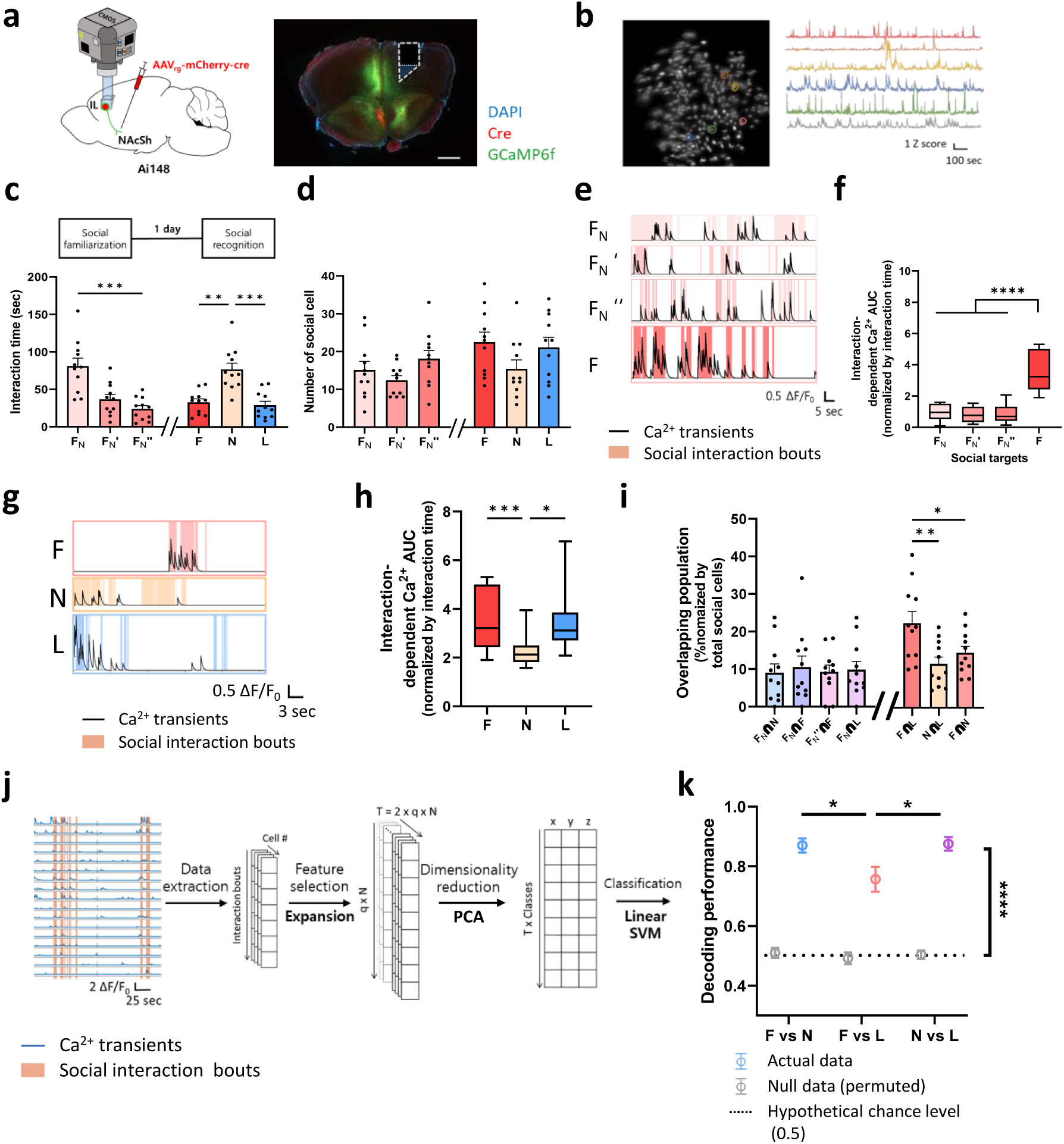
IL^→NAcSh^ neurons are activated by social interaction with familiar conspecifics during social memory retrieval. (a) Left: schematic diagram expressing GCaMP6f in IL^→NAcSh^ neurons in Ai148 mouse line for one photon Ca^2+^ imaging. Right: post hoc whole brain slice image showing GCaMP6f expression in IL and integrated lens traces for miniscope imaging (Scale bar: 500μm). (b) Example imaging plane of IL^→NAcSh^ neurons with pseudo-colored ROIs for Ca^2+^ analysis (left) and representative Ca^2+^ traces of the ROIs (right). (c) Subject mice showed significantly decreased interaction times over the trials during social familiarization. During the social recognition session, mice showed significantly higher interaction times with novel conspecifics compared to the familiar conspecifics or littermates. One-way ANOVA with repeated measures: n = 11; Effect of social target, F_3.325,33.25_ = 18.63, ****p < 0.0001. Tukey’s multiple comparison test: F_N_ versus F_N_’ F_10_ = 8.250, **p = 0.0017; F_N_ versus F_N_’’ F_10_ = 8.578, **p = 0.0024; F_N_’ versus F_N_’’ F_10_ = 2.509, p = 0.5199; F versus N F_10_ = 6.395, *p = 0.0103; F versus L F_10_ = 0.9692, p = 0.9796; N versus L F_10_ = 8.889, ***p = 0.0009. (d) Subject mice showed comparable number of social cells during the social familiarization/recognition task. Social familiarization session, one-way ANOVA with repeated measures: n = 11; effect of social target, F_1.750,17.50_ = 2.418, p = 0.1233. Social recognition session, one-way ANOVA with repeated measures: n = 11; effect of social target, F_1.380,13.80_ = 3.306, p = 0.0807. (e) A representative image of the neural activity of a social cell for familiar social target during the social familiarization/recognition task. Interaction bouts were labeled with colored columns. (f) Interaction-dependent Ca^2+^ AUC of the social cells for familiar social target significantly increased 24h after social familiarization. One-way ANOVA with repeated measures, F_1.596,15.96_ = 68.94, ****p<0.0001, Tukey’s multiple comparison test: F_N_ versus F F_10_ = 14.45, ****p<0.0001; F_N_’ versus F F_10_ = 14.51, ****p<0.0001; F_N_’’ versus F F_10_ = 11.44, ****p<0.0001. (g) Representative images showing the Ca^2+^ transients of the social cell during interaction with each corresponding social target: a familiar mouse (top), a novel mouse (center), and a littermate (bottom). Interaction bouts were labeled with colored columns. (h) Social interaction-dependent AUC from Ca^2+^ signal of social cells for known social targets showed a significantly larger AUC than novel conspecific social cell during each corresponding behavior session. One-way ANOVA with repeated measures, F_1.536,15.36_ = 10.41, **p = 0.0024, Tukey’s multiple comparison test: F versus N F_10_ = 7.692, ***p = 0.0008; F versus L F_10_ = 0.02946, p = 0.9998; N versus L F_10_ = 4.610, *p = 0.0213. (i) Social cells for known social targets exhibited significantly higher overlap compared to other population overlaps. One-way ANOVA with repeated measures (left): n = 11; effect of social target, F_1.982,19.82_ = 0.1608, p = 0.8507. One-way ANOVA with repeated measures (right): n = 11; effect of social target, F_1.317,13.17_ = 11.39, **p = 0.0030. Tukey’s multiple comparison test: F∩L versus N∩L q_10_ = 5.746, **p = 0.0059; F∩L versus F∩N q_10_ = 3.982, **p = 0.0443; N∩L versus F∩N q_10_ = 3.317, p = 0.0949. (j) Schematic diagram of Ca^2+^ data process which behavior-specific data being extracted, expanded, and dimensionally reduced (PCA) before being classified with a planar classifier (LSVM). A representative image of a planar classifier which decodes identity from vector representations of neural population has been shown. (k) LSVM classifier successfully decoded social targets, while decoding performance for distinguishing between familiar conspecifics (F) and littermate (L) was significantly lower than those for other comparisons. Two-way ANOVA: n = 10; Interaction between null data comparison and target comparison F_2,54_ = 2.797, p = 0.0698, Null data comparison F_1,54_ = 2699, ****p <0.0001, Comparison group F_2,54_ = 4.663, *p = 0.0135. One-way ANOVA with repeated measures: n = 10; Target comparison (between columns), F_1.714,15.42_ = 8.865, **p = 0.0037. Tukey’s multiple comparison test: F vs N versus N vs L q_9_ = 4.380, *p = 0.0311; F vs N versus N vs L q_9_ = 0.3107, p = 0.9738; F vs L versus N vs L q_9_ = 5.119, *p = 0.0139.

Social interaction-responsive cell population among the IL^→NAcSh^ neurons was defined by using a receiver operating characteristic (ROC) analysis and classified as ‘social cells (see “Methods” section for details)’^17^. 47.24% of total cells (558 out of 1181) were identified as social cells in at least one of the interaction sessions during two-days social familiarization/recognition task. The number of social cells were comparable between sessions (Fig. 2d). Interestingly, when the interaction-dependent Ca^2+^ transients’ area under curve (AUC) of familiar conspecific-social cells was analyzed to examine the impact of familiarization, there was a significant increase in AUC during the social recognition session compared to the familiarization session (Figs. 2e and 2f). Repeated interaction with a conspecific within the familiarization session did not increase either the number of social cells or overlapping between them (Figs. 2d and 2i). These data along with our optogenetic inhibition results during social memory encoding and retrieval suggest a delayed emergence of neural representation for familiar mice in the cortex 1 day after repeated social interaction.

Additionally, we found that the social cells for littermates and familiar conspecifics showed significantly larger interaction-dependent Ca^2+^ transient AUC compared to the social cells for novel conspecifics, which is consistent with our previous finding that IL^→NAcSh^ neurons show increased responses to familiar mice than novel mice (Figs. 2g and 2h)^26^. Then, we asked whether the familiar mice and littermates activate the same or overlapping subpopulations of IL^→NAcSh^ neurons during social recognition. We found that social cells for known conspecifics (F and L) showed significantly larger overlap (F∩L) than other comparison groups, though comparable size of social cells between sessions (Figs. 2d and 2i). These results led us to hypothesize that the IL^→NAcSh^ neural activities encode social familiarity, enabling them to more effectively distinguish between a novel mouse and a familiar mouse than between a familiar mouse and a littermate. To test this, we examined whether the population activity of IL^→NAcSh^ neurons could decode the identity of social targets at different levels of familiarity. We trained a 3-dimensional linear support vector machine (LSVM) decoding algorithm with two single variables after linear dimensionality reduction with principal component analysis (PCA) (Fig. 2j). IL^→NAcSh^ neuronal representations were able to accurately discriminate between mice regardless of their familiarity levels. However, the LSVM decoding performance for distinguishing between the two known social targets (F and L) was significantly lower compared to other comparison groups (Fig. 2k). In conjunction with the ROC-based identification of social interaction-dependent cell populations and their properties, these results imply that IL^→NAcSh^ neurons encode social memory of known social targets following social familiarization and subsequent memory consolidation.

### IL^→NAcSh^ neural activity represents social identity and familiarity

Recent studies have shown that neural ensembles in the PFC and hippocampus can simultaneously represent variables in an abstract format while still preserving their ability to decode other variables including social familiarity and identity^8,23^. A recent study showed that CA2 activity encodes both social identity and familiarity by analyzing how patterns of neural activity are arranged, known as the geometry of neural representations, in neural activity space^8^. Given the preferential activities of IL^→NAcSh^ neurons towards known social targets and the descent decoding performance for distinguishing between two known social targets, we questioned whether IL^→NAcSh^ neurons can simultaneously represent both social familiarity and identity of familiar conspecifics. We monitored Ca^2+^ activity of the IL^→NAcSh^ neurons while subject mice were consecutively exposed to either two littermates or two novel mice with the positions of the same target mice reversed between exposures (Figs. 3a and 3c). During the consecutive littermate recognition task, the subject mice interacted twice with each of two littermates alternatively, exhibiting a slight reduction in the interaction time in the second interaction with their littermates (Figs. 3a and 3b). A similar proportion of social cells was observed in each session (Supplementary Fig. 3a). Interestingly, the overlap between the social cells identified for the same littermates in different sessions was not higher than the overlap between social cells for different littermates, suggesting that the IL^→NAcSh^ neurons were not consistently activated by a specific littermate (Supplementary Fig. 3b). Nevertheless, the LSVM classifier trained either with social (identity) or spatial (position) information displayed a high decoding accuracy (Supplementary Figs. 3c–3e).

**Figure 3.**
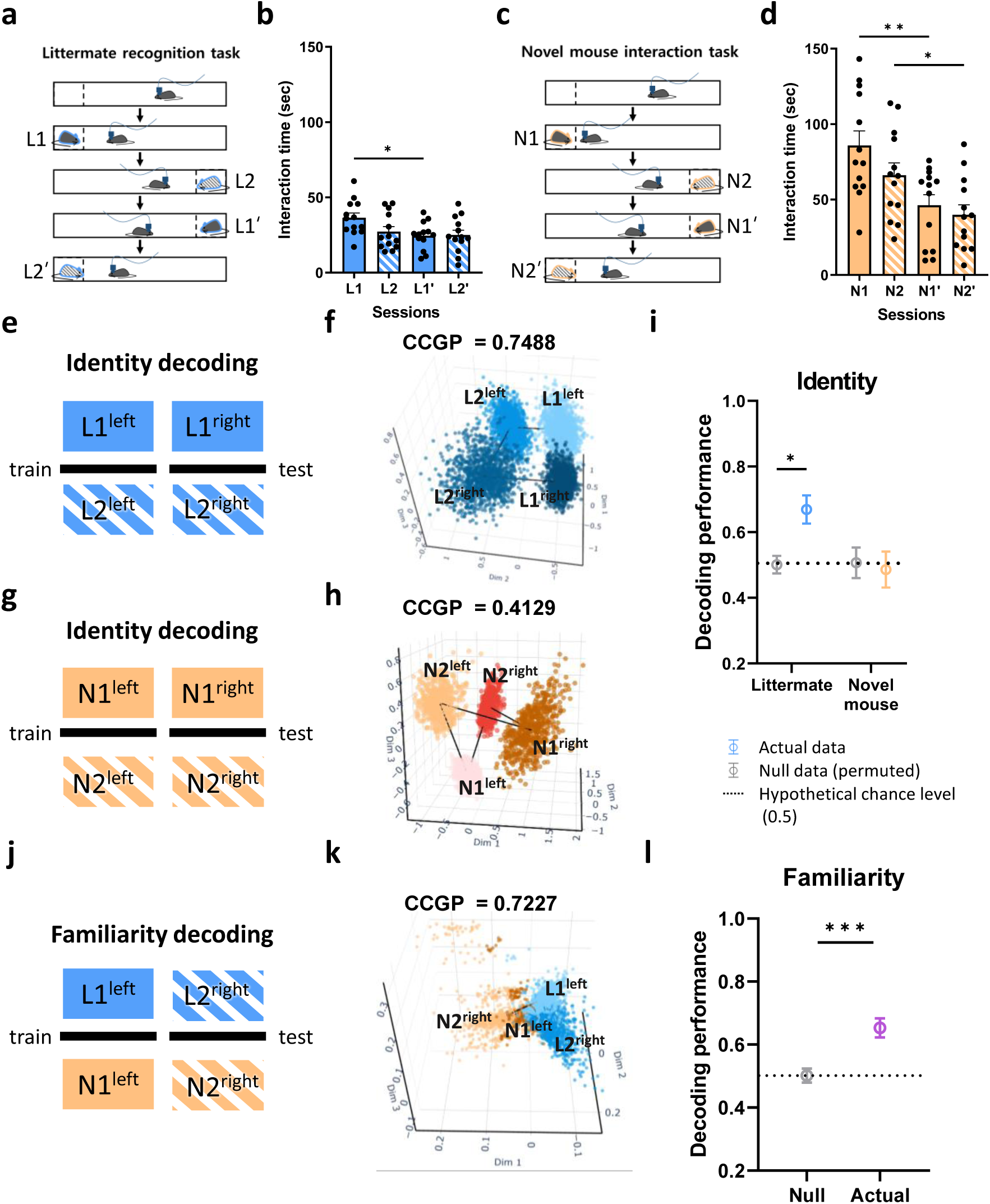
IL^→NAcSh^ neurons encode generalized social identity of familiar conspecifics. (a) Experimental scheme for consecutive littermate recognition task. Subject mice interacted with two littermates (L1 and L2) for 5 minutes each, twice with a 5-minute interval between each session. (b) Subject mice showed significantly decreased interaction times during interaction with their littermates. One-way ANOVA with repeated measures: n = 13 mice; F_2.342,28.10_ = 5.113, **p = 0.0097, Tukey’s multiple comparison test: L1 versus L1’ q_12_ = 5.110, *p = 0.0162. (c) Experimental scheme for consecutive novel mouse interaction task. Subject mice interacted with two novel conspecifics (N1 and N2) for 5 minutes each twice, with a 5-minute interval between each session. (d) Subject mice interacted with novel conspecifics, showing significantly decreased interaction times during interaction sessions. One-way ANOVA with repeated measures: n = 13 mice; F_2.562,30.74_ = 17.32, ****p < 0.0001, Tukey’s multiple comparison test: N1 versus N1’ q_12_ = 8.199, ***p = 0.0009, N2 versus N2’ q_12_ = 5.920, *p = 0.0124. (e) Schematic diagram for decoding identity using CCGP analysis from the consecutive littermate recognition task. (f) A representative image showing that IL^→NAcSh^ neural activity supports generalized identity decoding of littermate. (g) Schematic diagram for decoding identity using CCGP analysis from the consecutive novel mouse interaction task. (h) A representative image showing that IL^→NAcSh^ neural activity does not support generalized identity decoding of novel mice. (i) Social interaction-dependent neural activities from littermate interactions showed significantly higher CCGP decoding performance than those from null model, while those from novel conspecific interactions exhibited comparable CCGP decoding performance to those from null model. Two-way ANOVA with Sidak’s multiple comparison test: n = 11, F_1,82_ = 4.523, *p = 0.0365, Littermate; Null-Actual: t_82_ = 2.649, *p = 0.0192, Novel mouse; Null-Actual: t_82_ = 0.3314, p = 0.9330. (j) Schematic diagram for decoding familiarity with CCGP analysis by comparing social interaction-dependent IL^→NAcSh^ neural activity from littermate and novel mouse interactions. (k) A representative image of social interaction-dependent IL^→NAcSh^ neural activity from littermates and novel mouse for social familiarity CCGP analysis. (l) Social familiarity CCGP analysis by comparing social interaction-dependent IL^→NAcSh^ neural activity from littermate and novel mouse interaction exhibited significantly high decoding performance compared to that from null data. Paired t-test: n = 9 mice, CCGP; t_8_ = 6.500. ***p = 0.0002.

During the consecutive novel mouse interaction task, the subject mice alternately interacted twice with two novel mice (Fig. 3c). Interaction times with the novel mice were significantly higher than those with littermates during the first encounter but then decreased in the subsequent interactions to levels comparable to those with littermates (Figs. 3b and 3d). The number of social cells for novel conspecifics remained constant across sessions (Supplementary Fig. 3f). The overlap between social cells for different or same novel mice was comparable, suggesting these neurons might not distinguish individual novel mice, like social cells for littermates (Supplementary Figs. 3b and 3g). Consistent with the observation in the social familiarization/recognition task, Ca^2+^ responses of the social cells for novel mice were substantially smaller than those of social cells during interactions with littermates (Supplementary Fig. 3h). Despite the overall lower Ca^2+^ activity of IL^→NAcSh^ social cells during interactions with novel mice, both social identity and position of novel mice could be successfully decoded by the population activity of IL^→NAcSh^ neurons (Supplementary Figs. 3i-3k).

As we found that social and non-social (spatial) features of social targets can be decoded from the activity of the IL^→NAcSh^ neurons by using a linear classifier, we then explored whether social identity could be represented independently from the position in the geometry supporting generalization. To answer this question, a modified linear SVM classifier called cross-condition generalization performance (CCGP) was used to train and determine a conceptual hyperplane that can separate population of vectors illustrating different neural representations that are effectively generalized to predict^8,23^. We measured identity CCGP by training the linear classifier to decode the identity of two littermates on the left side (L1^left^ and L2^left^) and tested on data from littermates on right side (L1^right^ and L2^right^) (Fig. 3e). We also measured position CCGP. We found high CCGP performance for identity of littermates, but not for their spatial information, suggesting that the IL^→NAcSh^ neural activity supports generalized decoding of identity, but not position of littermates (Figs. 3f, 3i and Supplementary Figs. 4a, 4b, and 4e). Interestingly, data from novel mouse interactions revealed CCGP performance as low as chance level for both identity and position (Figs. 3g-i and Supplementary Figs. 4c-4e).

Next, we directly investigated the effect of familiarity of social targets on IL^→NAcSh^ neuronal representations, by comparing the population patterns from two behavioral paradigms – littermate recognition task and novel mouse interaction task. Cells from each behavior paradigms were registered (Supplementary Figs. 5a-d). The CCGP classifier was trained using data from interactions with littermate 1 and novel mouse 1 on the left side (L1^left^ and N1^left^) and tested on data from littermate 2 and novel mouse 2 on the right side (L2^right^ and N2^right^). The reversed positional configuration was also applied, and the results were averaged to account for side-specific bias (Fig. 3j). Considering different social identities and positions of target mice, the CCGP classifier specifically allows determining the impact of familiarity beyond identity. We found that IL^→NAcSh^ neural activity showed significantly high decoding performance depending on the familiarity of social targets with different identities, but not the spatial location of social targets (Figs. 3j–3l, and Supplementary Figs. 4f-h). These data suggest that IL^→NAcSh^ neural activities can not only represent generalized social identity of littermates, but also support the representation of familiarity. Of note, LSVM decoder failed to distinguish the calcium activity patterns from the habituation sessions of littermate recognition and novel mouse interaction task, showing that the decoding performance does not depend on time differences between two tasks (Supplementary Fig. 5e).

### Inactivation of IL^→NAcSh^ neurons activated by a familiar mouse impairs social recognition

To investigate whether selective inhibition of IL^→NAcSh^ social cells activated by a familiar conspecific, hence disrupting the neuronal representation of a single known social target, is sufficient to disrupt the recognition of other conspecifics, we used a genetic technique known as targeted recombination in active population (TRAP)^27^. We injected the fosCreERT2 virus in the IL cortex and retrograde Cre-dependent virus expressing hM4Di or mCherry in the NAcSh of wild-type mice (Fig. 4a). IL^→NAcSh^ neurons activated by a familiar conspecific were labelled (TRAPed) and their activities were subsequently manipulated during the social recognition session (Fig. 4b). Subject mice were exposed to a novel conspecific for familiarization on day 1 and re-exposed to the familiarized mouse for TRAP tagging on day 2 (Fig. 4c). One week later (day 9) when either hM4Di-mCherry (hereafter hM4Di) or mCherry was expressed in TRAPed neurons, the subject mice were tested in the social familiarization/recognition task, in which the subject mice were familiarized with another novel mouse different from the one interacted with on day 1 and 2 (Figs. 4c and 4d). Both hM4Di and mCherry-expressing groups displayed similar decreases in interaction times during familiarization with a novel conspecific on day 9 regardless of CNO injection (Fig. 4d, Supplementary Figs. 6a and 6d). However, inactivating TRAPed IL^→NAcSh^ neurons by injecting CNO before the social recognition session significantly impaired social recognition on day 10 compared to the control groups including mCherry-expressing mice with or without CNO injection and hM4Di-expressing mice without CNO injection (Fig. 4d, Supplementary Figs. 6b, 6c, 6e, and 6f). Importantly, interaction time not only with a familiarized mouse, but also with a littermate was significantly increased in CNO-injected hM4Di-expressing mice (Fig. 4d and Supplementary Fig. 6f). Basal locomotor activity and total interaction times were not affected by inactivating TRAPed IL^→NAcSh^ neurons (Supplementary Figs. 6g and 6h). These data show that inactivation of a single familiar conspecific-dependent IL^→NAcSh^ neurons is sufficient to impair social recognition of other known social targets, demonstrating that IL^→NAcSh^ neuronal activity supports social familiarity.

**Figure 4.**
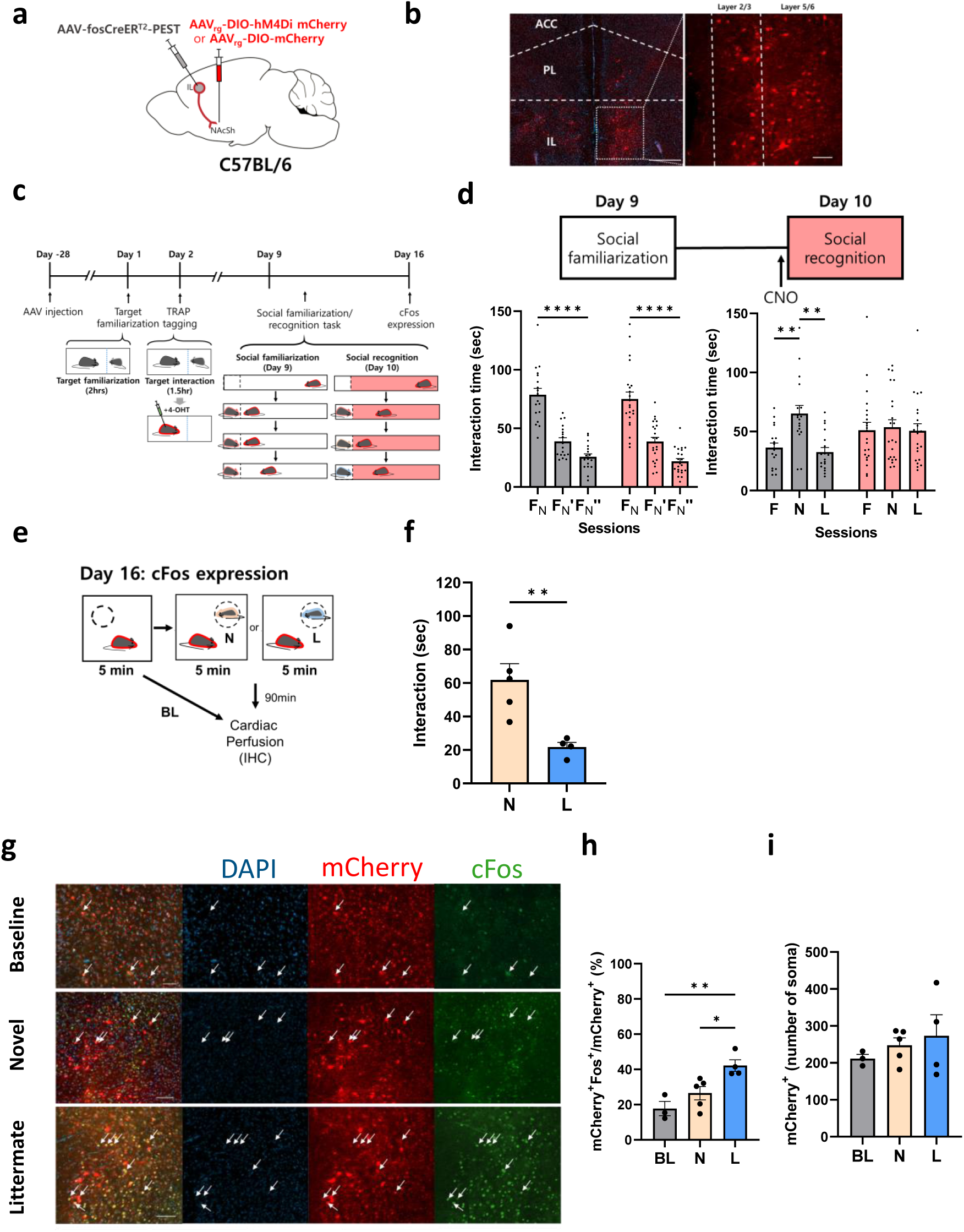
Inactivation of IL^→NAcSh^ neurons activated by a familiar mouse impairs social recognition. (a) Schematic diagram of TRAP tagging in IL^→NAcSh^ neurons. (b) Left: a representative image of mCherry expression (TRAP signal) in the mPFC after exposed to a familiar conspecific (Scale bar: 500μm). Right: mCherry (or hM4Di-mCherry) was expressed in the IL (Scale bar: 100μm). ACC, anterior cingulate cortex; PL, prelimbic cortex; IL, infralimbic cortex (c) Schematic diagram of labelling familiar conspecific-dependent neurons using TRAP. Day 1: A subject mouse was familiarized with a social target by 2 hrs of social interaction. Day 2: The subject mouse interacted with previously familiarized social target followed by tamoxifen (4-OHT) injection to tag activated cells. Day 9-10: The subject mouse underwent the social familiarization/recognition task with independent novel mice. (d) During the social familiarization/recognition task, CNO or saline (as a vehicle) was injected to the subject mice 40 minutes prior to the social recognition. Both mCherry- and hM4Di-expressing-mice showed significantly decreased interaction times during the social familiarization (left). Two-way ANOVA with repeated measures: mCherry CNO, n = 18 mice; hM4Di CNO, n = 22 mice; interaction between session and group, F_2,76_ = 0.1825, p = 0.8336. Tukey’s multiple comparison test: mCherry CNO, F_N_ versus F_N_’ q_76_ = 11.02, ****p < 0.0001; mCherry CNO, F_N_ versus F_N_’’ q_76_ = 14.66, ****p < 0.0001; mCherry CNO, F_N_’ versus F_N_’’ q_76_ = 3.639, *p = 0.0319; hM4Di CNO, F_N_ versus F_N_’ q_76_ = 11.12, ****p < 0.0001; hM4Di CNO, F_N_ versus F_N_’’ q_76_ = 16.28, ****p < 0.0001; hM4Di CNO, F_N_’ versus F_N_’’ q_76_ = 5.159, **p = 0.0014. CNO-injected mCherry-expressing mice showed a comparable interaction time with different conspecifics, while CNO-injected hM4Di-expressing mice showed a significantly longer interaction times with novel conspecifics than familiar conspecifics or littermates (right). Two-way ANOVA with repeated measures: mCherry CNO, n = 18 mice; hM4Di CNO, n = 22 mice; interaction between target effect and CNO effect, F_2,76_ = 5.170, **p = 0.0079. Tukey’s multiple comparison test: mCherry CNO, F versus N q_17_ = 4.888, **p = 0.0080; mCherry CNO, F versus L q_17_ = 1.256, p = 0.6549; mCherry CNO, N versus L q_17_ = 5.552, **p = 0.0030; hM4Di CNO, F versus N q_21_ = 0.4431, p = 0.9475; hM4Di CNO, F versus L q_21_ = 0.1688, p = 0.9922; hM4Di CNO, N versus L q_21_ = 0.5686, p = 0.9151. (e) Schematic diagram of cFos induction by novel conspecifics or littermate interaction in TRAP-labeled (TRAPed) mice on day 16. BL, baseline; N, novel mouse; L, littermate. (f) Subject mice showed significantly longer interaction times with novel conspecifics than littermates during cFos induction. Unpaired t-test: TRAPed mice interacted with littermate, n = 4; TRAP mice interaction with novel conspecific, n = 5, t_7_ = 3.571, **p = 0.0091. (g) Representative images of the IL exhibiting TRAPed neurons (mCherry) and cFos (green), in the basal state or after induction by novel social targets or littermates (Scale bar: 50 μm). (h) TRAPed neurons in mice interacted with littermates showed a significantly more overlap with cFos compared to those in mice without social interaction or mice interacting with novel conspecifics. One-way ANOVA: Baseline, n = 3 mice; Novel, n = 5 mice, Littermate, n = 4 mice; F_2,9_ = 9.531, **p = 0.0060. Tukey’s multiple comparison test: Baseline versus Novel q_9_ = 2.233, p = 0.3027; Baseline versus Littermate q_9_ = 5.950, **p = 0.0058; Littermate versus Novel q_9_ = 4.343, *p = 0.0324. (i) Mice without any social interaction, mice interacting with littermates, and mice interacting with novel conspecifics showed a comparable number of TRAPed neurons. One-way ANOVA: Baseline, n = 3 mice; Novel, n = 5 mice; Littermate, n = 4 mice, F_2,9_ = 0.6061, p = 0.5663. Tukey’s multiple comparison test: Baseline versus Novel q_9_ = 0.9472, p = 0.7862; Baseline versus Littermate q_9_ = 1.557, p = 0.5372; Littermate versus Novel q_9_ = 0.7414, p = 0.8616. BL: Baseline, N: Novel social target, L: Littermate.

To further investigate whether the TRAPed IL^→NAcSh^ neuronal population represents social familiarity, we examined whether these neurons are reactivated during interactions with another familiar conspecifics. The TRAPed mice were allowed to interact with either a novel conspecific or a littermate, followed by analyzing cFos expression (day 15-16, Figs. 4c and 4e). The TRAPed mice showed a strong preference towards novel conspecifics over their littermates (Fig. 4f). We found that TRAPed mice interacted with a littermate showed a significantly higher percentage of co-labeled cFos with TRAPed mCherry^+^ neurons compared to mice interacted with a novel mouse or baseline control without any social interactions (Figs. 4g and 4h). The numbers of TRAP-labeled neurons (mCherry^+^) were comparable among groups (Fig. 4i). The observed higher overlap between TRAPed neurons and littermate-induced cFos^+^ neurons is consistent with the high overlap of social cells between known social targets in the Ca^2+^ imaging experiments (Fig. 2i). These results illustrate that IL^→NAcSh^ neurons encode social memory with a certain level of overlap, which might support social familiarity.

### vCA1 to IL projections are necessary for social memory consolidation

Given that the hippocampus plays crucial roles in multiple phases of social memory including memory acquisition and the hippocampal-cortical interaction is critical for episodic memory, we hypothesized that the IL might interact with the hippocampus to process social memory. Among subregions of the hippocampus engaged in social memory, the IL cortex receives direct inputs from vCA1^28^ . We investigated the temporal window that requires the activity of IL-projecting vCA1 (vCA1^→IL^) neurons in social recognition with either optogenetic or chemogenetic manipulation. We expressed NpHR or YFP in the vCA1 and optic fibers were bilaterally targeted in the IL to selectively inactivate vCA1^→IL^ neurons (Fig. 5a). Illuminating 594 nm laser during social familiarization did not affect social recognition in either NpHR- or YFP-expressing in the social recognition session (Fig. 5b and Supplementary Fig. 7a).

**Figure 5.**
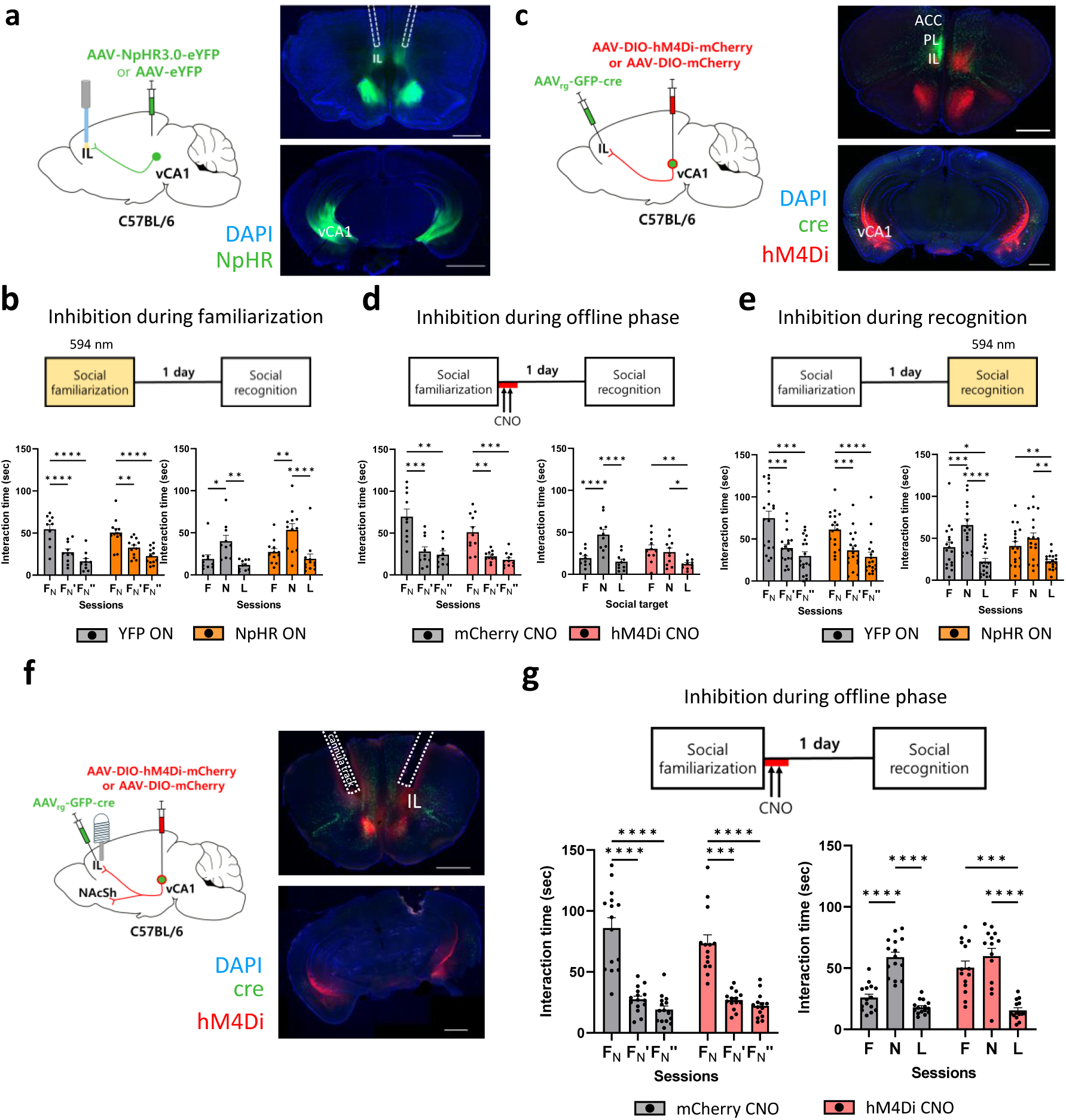
Inactivation of vCA1^→IL^ neuron and its terminals during the offline consolidation and retrieval impairs social recognition. (a) Left: schematic diagram of optogenetic inactivation of vCA1^→IL^ neurons. NpHR3.0 or eYFP was expressed in vCA1 neurons while two 200 μm optic fibers were bilaterally implanted in the IL. Right: representative image of mPFC (top, Scale bar: 1000 μm) and vCA1 (bottom, Scale bar: 2000 μm) expressing NpHR in neuronal terminal and soma, respectively. (b) Left: both YFP and NpHR-expressing mice showed significantly decreased interaction times during social familiarization. Two-way ANOVA with repeated measures: YFP, n = 10 mice;1 NpHR, n = 12 mice; interaction between session and NpHR effect, F_2, 40_ = 1.396, p = 0.2593. Tukey’s multiple comparison test: YFP, F_N_ versus F_N_’ q_40_ = 7.599, ****p < 0.0001; YFP, F_N_ versus F_N_’’ q_40_ = 10.60, ****p < 0.0001; YFP, F_N_’ versus F_N_’’ q_40_ = 3.003, p = 0.0.0978; NpHR, F_N_ versus F_N_’ q_40_ = 5.423, **p = 0.0012; NpHR, F_N_ versus F_N_’’ q_40_ = 8.459, ****p < 0.0001; NpHR, F_N_’ versus F_N_’’ q_40_ = 3.036, p = 0.0932. Right: both YFP and NpHR mice showed significantly longer interaction times with novel conspecifics than familiar conspecifics or littermates. Two-way ANOVA with repeated measures: YFP, n = 10 mice; NpHR, n = 12 mice; interaction between target effect and familiarization-laser effect, F_2, 40_ = 0.2207, p = 0.8029. Tukey’s multiple comparison test: YFP, F versus N q_40_ = 4.175, *p = 0.0142; YFP, F versus L q_40_ = 1.448, p = 0.5664; YFP, N versus L q_40_ = 5.623, ***p = 0.0008; NpHR, F versus N q_40_ = 5.674, ***p =0.0007; NpHR, F versus L q_40_ = 1.776, p = 0.4281; NpHR, N versus L q_40_ = 7.450, ****p < 0.0001. (c) Left: schematic diagram of expressing hM4Di or mCherry in vCA1^→IL^ neurons by injecting a Cre-dependent hM4Di or mCherry virus in the vCA1 and retrograde eGFP-Cre virus in the IL of mice. Right: representative image of mPFC (top) and vCA1 (bottom) expressing hM4Di in neuronal terminal and soma, respectively. (d) Left: both mCherry and hM4Di-expressing mice showed significantly decreased interaction times during the social familiarization. Two-way ANOVA with repeated measures: mCherry, n = 10 mice; hM4Di, n = 11 mice; interaction between session and offline-CNO effect, F_2, 38_ = 1.384, p = 0.2630. Tukey’s multiple comparison test: mCherry, F_N_ versus F_N_’ q_9_ = 10.11, ***p = 0.0001; mCherry, F_N_ versus F_N_’’ q_9_ = 7.765, **p = 0.0010; mCherry, F_N_’ versus F_N_’’ q_9_ = 1.011, p = 0.7612; hM4Di, F_N_ versus F_N_’ q_10_ = 5.521, **p = 0.0075; hM4Di, F_N_ versus F_N_’’ q_10_ = 7.421, ***p = 0.0010; hM4Di, F_N_’ versus F_N_’’ q_10_ = 1.844, p = 0.4245. Right: hM4Di mice showed comparable interaction times between familiar and novel conspecifics but significantly shorter interaction times with littermates, while mCherry mice showed significantly longer interaction times towards novel conspecifics than familiar conspecifics or littermates. Two-way ANOVA with repeated measures: mCherry, n = 10 mice; hM4Di, n = 11 mice; interaction between target effect and offline-CNO effect, F_2, 57_ = 7.197, **p = 0.0016. Tukey’s multiple comparison test: mCherry, F versus N q_57_ = 6.526, ****p < 0.0001; mCherry, F versus L q_57_ = 0.9873, p = 0.7655; mCherry, N versus L q_57_ = 7.513, ****p < 0.0001; hM4Di, F versus N q_57_ = 0.9119, p = 0.7960; hM4Di, F versus L q_57_ = 4.439, **p = 0.0075; hM4Di, N versus L q_27_ = 3.527, *p = 0.0406. (e) Left: both YFP and NpHR-expressing mice showed significantly decreased interaction times during the social familiarization. Two-way ANOVA with repeated measures: YFP, n = 18 mice; NpHR, n = 18 mice; interaction between session and recognition-laser effect, F_2, 68_ = 1.165, p = 0.3179. Tukey’s multiple comparison test: YFP, F_N_ versus F_N_’ q_17_ = 6.719, ***p = 0.0005; YFP, F_N_ versus F_N_’’ q_17_ = 7.471, ***p = 0.0002; YFP, F_N_’ versus F_N_’’ q_17_ = 2.183, p = 0.2964; NpHR, F_N_ versus F_N_’ q_17_ = 6.441, ***p = 0.008; NpHR, F_N_ versus F_N_’’ q_17_ = 10.70, ****p < 0.0001; NpHR, F_N_’ versus F_N_’’ q_17_ = 2.289, p = 0.2652. Right: NpHR mice showed comparable interaction times between familiar and novel conspecifics but significantly shorter interaction times with littermates. YFP mice showed significantly longer interaction times with novel conspecifics than familiar conspecifics or littermates. Two-way ANOVA with repeated measures: YFP, n = 18 mice; NpHR, n = 18 mice; interaction between target effect and recognition-laser effect, F_2, 68_ = 2.402, p = 0.0982. Tukey’s multiple comparison test: YFP, F versus N q_17_ = 6.491, ***p = 0.0007; YFP, F versus L q_17_ = 4.500, *p = 0.0143; YFP, N versus L q_17_ = 9.002, ****p < 0.0005; NpHR, F versus N q_17_ = 1.822, p =0.4205; NpHR, F versus L q_17_ = 5.050, **p = 0.0063; NpHR, N versus L q_17_ = 5.659, **p = 0.0025. (f) Left: schematic diagram of expressing hM4Di or mCherry specifically in IL^→NAcSh^ neurons. Infusion cannulas were bilaterally implanted targeting the IL cortex (Scale bar: 1000 μm). Right: representative image of coronal sections with the IL (top) and the vCA1 (bottom), respectively (Scale bar: 1000 μm). (g) Top: experimental scheme for the social familiarization/recognition task with chemogenetic inactivation during offline phase. CNO (200 nL, 1 mM) was given bilaterally to subject mice after the social familiarization task and once again two hours later. Bottom: both mCherry and hM4Di mice showed significantly decreased interaction times during the social familiarization task (left). Two-way ANOVA with repeated measures: mCherry, n = 15 mice; hM4Di, n = 14 mice; interaction between session and offline - CNO effect, F_2,24_ = 1.669, p = 0.1979. Tukey’s multiple comparison test: mCherry, F_N_ versus F_N_’ q_14_ = 11.53, ****p<0.0001; mCherry, F_N_ versus F_N_’’ q_14_ = 12.52, ****p<0.0001; mCherry, F_N_’ versus F_N_’’ q_14_ = 5.218, **p = 0.0064; hM4Di, F_N_ versus F_N_’ q_13_ = 8.536, ***p = 0.0001; hM4Di, F_N_ versus F_N_’’ q_13_ = 9.087, ****p<0.0001; hM4Di, F_N_’ versus F_N_’’ q_13_ = 2.253, p = 0.2831. hM4Di mice showed comparable interaction times between familiar and novel conspecifics while mCherry mice showed significantly longer interaction times towards novel conspecifics than familiar conspecifics or littermates during the social recognition task (right). Two-way ANOVA with repeated measures: mCherry, n = 15 mice; hM4Di, n = 14 mice; interaction between session and offline - CNO effect, F_2,54_ = 9.197, ***p = 0.0004. Tukey’s multiple comparison test: mCherry, F versus N q_14_ = 6.942, ****p<0.0001; mCherry, F versus L q_14_ = 3.389, *p = 0.0132; mCherry, N versus L q_14_ = 11.44, ****p<0.0001; hM4Di, F versus N q_13_ = 1.592, p = 0.3538; hM4Di, F versus L q_13_ = 6.173, ***p = 0.0001; hM4Di, N versus L q_13_ = 7.493, ****p < 0.0001.

For chemogenetic inhibition of vCA1^→IL^ neurons during offline memory consolidation phase, we expressed hM4Di in vCA1^→IL^ neurons (Fig. 5c). When these neurons were chemogenetically inactivated by CNO injection after the social familiarization, hM4Di-expressing mice showed deficits in social recognition: they showed comparable interaction times between familiar and novel conspecifics during the social recognition session (Fig. 5d). Interestingly, their interaction times with littermates still remained low as controls (Fig. 5d and Supplementary Fig. 7b). In addition, optogenetic inhibition of vCA1^→IL^ neurons during retrieval phase also prevented NpHR-expressing mice from distinguishing between familiar and novel conspecifics during the social recognition session (Fig. 5e and Supplementary Fig. 7c). This could be attributed to either incomplete memory transfer from the hippocampus to the cortex rather than a block in memory recall, as the subject mice still showed lower interaction times with littermates (Fig. 5e). Both YFP and NpHR-expressing mice showed intact locomotor activity and place preference regardless of the presence of 594 nm laser (Supplementary Figs. 8a and 8b). Chemogenetic inactivation of vCA1^→IL^ neurons did not affect novel object recognition, suggesting the specific engagement of the projection in social context (Supplementary Fig. 8c).

As previously reported^29^, we observed that vCA1^→IL^ neurons have collateral projections to the NAcSh (Figs. 5a and 5c, Supplementary Figs. 9). This raised a concern that the social memory deficits may arise from blocking the activity of NAcSh-projecting vCA1 neurons in the previous experiment (Fig. 5d). To selectively inactivate vCA1^→IL^ neurons, we expressed hM4Di in IL-projecting vCA1 neurons and locally infused CNO into the IL (Fig. 5f). While both mCherry and hM4Di-expressing mice showed normal behaviors during the social familiarization session, hM4Di-expressing mice injected with CNO during the offline phase exhibited comparable interaction times with familiar conspecifics to those with novel mice, which mainly due to increased investigating times towards familiar conspecifics (Fig. 5g, Supplementary Figs. 7d and 7e). Consistent with systemic CNO injection result (Fig. 5d), their interaction times with littermates were also maintained as low as controls. These data imply that the activity of IL-projecting vCA1 neurons during offline phase is critical for consolidation of memory for newly familiarized mice.

### Inactivation of vCA1^→IL^ neurons during offline phase impairs social recognition and social representation in IL^→NAcSh^ neurons

As the inactivation of hippocampus during offline period prevents consolidation of spatial or contextual memories^30^, we examined the impact of inactivating vCA1^→IL^ neurons during offline phase on the neural representation of social information in the IL cortex. First, we used two recombination systems (Cre-Lox and fLp-FRT system) to selectively express GCaMP protein in IL^→NAcSh^ neurons and hM4Di or mCherry (control) in vCA1^→IL^ neurons (Fig. 6a). Then, we recorded the Ca^2+^ activity of IL^→NAcSh^ neurons during an extended version of the social familiarization/recognition task (Fig. 6b and Supplementary Figs. 10a-d). In this task, a subject mouse was familiarized with two social targets during the social familiarization session and then interacted with those two familiarized mice, two novel conspecifics, and two littermates on the next day to test its social recognition. In this extended social familiarization/recognition task, mCherry-expressing control mice injected with CNO showed significantly longer interaction times with two novel mice compared to those either with two familiar mice or littermates (Fig. 6c). Consistent with the previous results (Figs. 5d and 5g), hM4Di-expressing mice injected with CNO during offline phase failed to distinguish familiar mice from novel mice (Fig. 6d). Importantly, these mice still showed significantly shorter interaction times with their littermates compared to those with familiar or novel mice, further demonstrating that vCA1 inactivation led subject mice to fail to recognize familiarized conspecifics as familiar and to show increased interaction with them (Fig. 6d).

**Figure 6.**
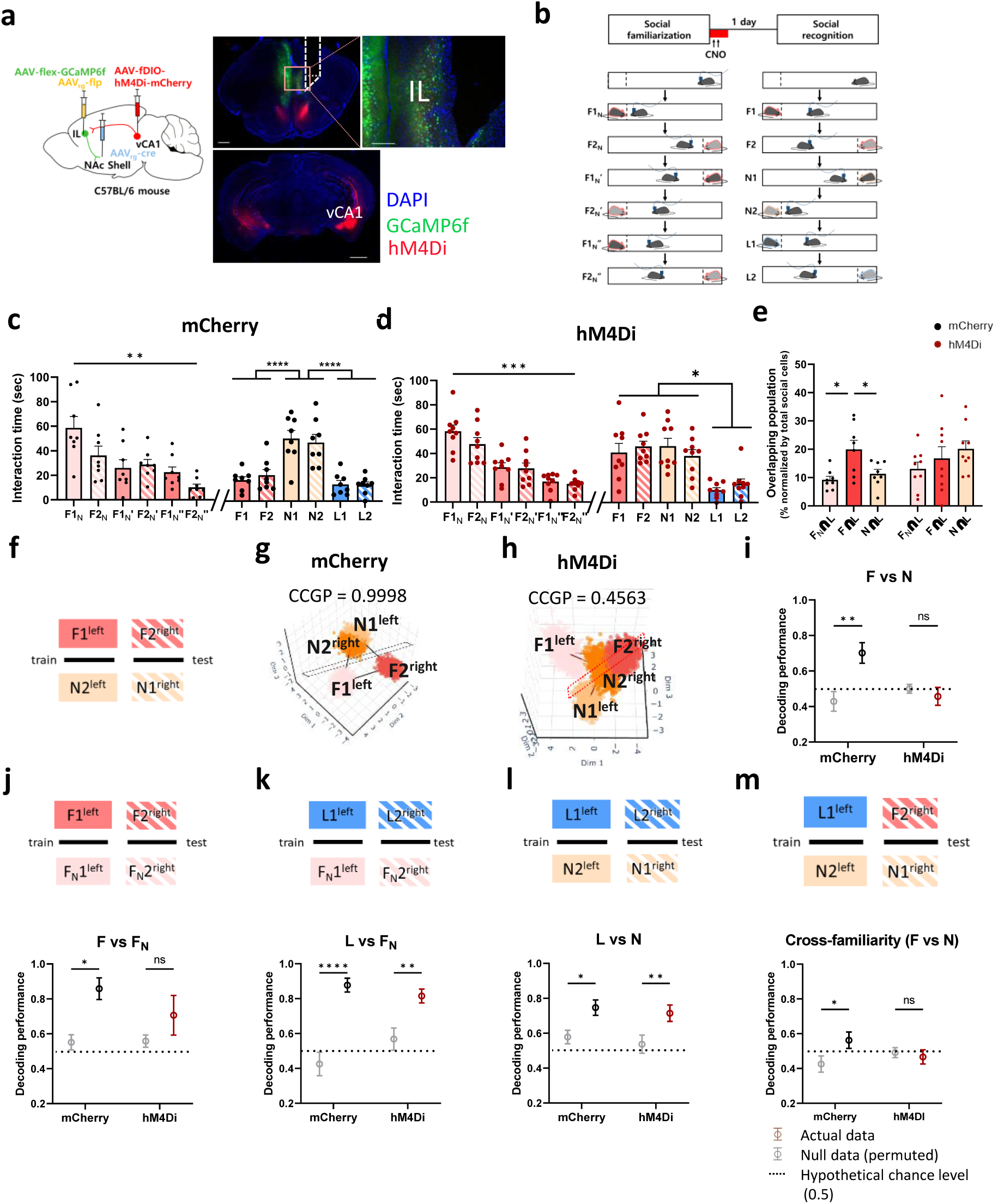
vCA1 inactivation during the offline phase disrupts social memory representations in IL^→NAcSh^ neurons. (a) Left: schematic diagram of selective expression of GCaMP6f in IL^→NAcSh^ neurons and hM4Di in vCA1^→IL^ neurons. Right top: representative image of IL expressing GCaMP6f. Scale bar: 500 µm, 100 µm. Right bottom: representative image of vCA1 expressing hM4Di. Scale bar: 1000 µm. (b) Schematic diagram of the extended social familiarization/recognition task while inactivating vCA1^→IL^ neurons during the offline phase. Each subject mouse was familiarized with two social targets (F_N_1 and F_N_2) during the social familiarization task. The subject interacted with two familiar conspecifics (F1 and F2), two novel conspecifics (N1 and N2), and two littermates (L1 and L2). CNO was systemically injected to subject mice immediately after the social familiarization session and once again two hours later. (c) Left: mCherry-expressing mice showed significantly decreased interaction times during the social familiarization session. One-way ANOVA with repeated measures: mCherry, n = 8 mice; F_2.800,19.60_ = 9.807, ***p = 0.0004. Tukey’s multiple comparison test: F1_N_ versus F 1_N_’’ q_7_ = 6.957, *p = 0.0137; F1_N_ versus F2_N_’’ q_7_ = 8.422, **p = 0.0047; F2_N_’ versus F2_N_’’ q_7_ = 5.899, *p = 0.0318. Right: during the social recognition session, mCherry-expressing mice exhibited significantly longer interaction times with novel mice than familiar mice or littermates. One-way ANOVA with repeated measures: mCherry, n = 8 mice; F_2.304,16.13_ = 21.59, ****p < 0.0001. Tukey’s multiple comparison test: F1 versus N1 q_7_ = 7.097, *p = 0.0123; F1 versus F2 q_7_ = 7.023, *p = 0.0130; F2 versus N1 q_7_ = 5.747, *p = 0.0361; F2 versus N2 q_7_ = 7.936, **p = 0.0066; F2 versus L1 q_7_ = 5.944, *p = 0.0307; N1 versus L1 q_7_ = 7.809, **p = 0.0073; N1 versus L2 q_7_ = 9.451, **p = 0.0024; N2 versus L1 q_7_ = 9.768, **p = 0.0020; N2 versus L1 q_7_ = 8.796, **p = 0.0037. (d) Left: hM4Di-expressing mice showed significantly decreased interaction times during the social familiarization session. One-way ANOVA with repeated measures: hM4Di, n = 9 mice; F_2.929,23.43_ = 17.23, ****p < 0.0001. Tukey’s multiple comparison test: F_N_1 versus F_N_1’ q_8_ = 7.741, **p = 0.0052; F_N_1 versus F_N_1’’ q_8_ = 11.12, ***p = 0.0005; F_N_1 versus F_N_2’’ q_8_ = 11.92, ***p = 0.0003; F_N_2 versus F_N_1’’ q_8_ = 8.849, **p = 0.0022; F_N_2 versus F_N_2’’ q_8_ = 7.825, **p = 0.0049. Right: during the social recognition task, hM4Di-expressing mice exhibited comparable interaction times between familiar and novel mice but significantly lower interaction times towards littermates. One-way ANOVA with repeated measures: hM4Di, n = 9 mice; F_3.259,26.07_ = 10.61, ****p < 0.0001. Tukey’s multiple comparison test: F1 versus L1 q_8_ = 6.254, *p = 0.0185; F1 versus L2 q_8_ = 4.382, p = 0.1047; F2 versus L1 q_8_ = 12.80, ***p = 0.0002; F2 versus L2 q_8_ = 6.579, *p = 0.0139; N1 versus L1 q_8_ = 7.644, **p = 0.0057; N1 versus L2 q_8_ = 5.292, *p = 0.0445; N2 versus L1 q_8_ = 9.965, ***p = 0.0010; N2 versus L2 q_8_ = 3843, p = 0.1732. (e) mCherry-expressing mice exhibited significantly increased overlapping social cells after familiarization, which was not observed in hM4Di-expressing mice. Two-way ANOVA with repeated measures, mCherry, n = 8, hM4Di, n = 9, interaction F_2, 30_ = 3.384, *p = 0.0473; mCherry, F_N_∩L/ Social versus F∩L/ Social t_30_ = 3.187, *p = 0.0100, F_N_∩L/ Social versus N∩L/ Social t_30_ = 0.6279, p = 0.8993, F∩L/ Social versus N∩L/ Social t_30_ = 2.559, *p = 0.0466; hM4Di, F_N_∩L/ Social versus F∩L/ Social t_30_ = 1.209, p = 0.5811, F_N_∩L/ Social versus N∩L/ Social t_30_ = 2.229, p = 0.0971, F∩L/ Social versus N∩L/ Social t_30_ = 1.060, p = 0.6533. (f) Schematic diagram for decoding familiarity with CCGP analysis. (g) A representative geometry image of mCherry-expressing mice exhibiting high CCGP performance for social familiarity. (h) A representative geometry image of hM4Di-expressing mice showing low CCGP performances for social familiarity as the chance level. (i) While mCherry-expressing mice showed significantly higher CCGP performance scores for the familiarity when comparing between familiar and novel social targets, hM4Di-expressing mice showed a chance level of CCGP performance compared to those of null models. Two-way ANOVA with repeated measures: mCherry, n = 8 mice; hM4Di, n = 9 mice; interaction between hM4Di effect and null model effect, F_1,15_ = 9.343, **p = 0.0080. Sidak’s multiple comparison test: mCherry, Null versus Actual t_15_ = 3.636, **p = 0.0049; hM4Di, Null versus Actual t_15_ = 0.5997, p = 0.8043. (j) Top: schematic diagram representing familiarity CCGP decoding between familiar conspecifics and unfamiliarized conspecifics. Bottom: while mCherry-expressing mice showed significantly higher CCGP performances than CCGP performances from the null models, hM4Di-expressing mice showed comparable CCGP performances to the null models. Two-way ANOVA with stacked matching: mCherry n = 8 mice, hM4Di n = 9 mice; interaction between hM4Di effect and null model effect, F_1, 15_ = 0.9767, p = 0.3387. Sidak’s multiple comparison test: mCherry, Null versus Actual t_15_ = 2.623, *p = 0.0381, hM4Di, Null versus Actual t_15_ = 1.341, p = 0.3598. (k) Top: schematic diagram representing familiarity CCGP decoding performance between littermates and unfamiliarized conspecifics. Bottom: both mCherry- and hM4Di-expressing mice showed significantly higher CCGP performances than CCGP performances from the null models. Two-way ANOVA with stacked matching: mCherry n = 8 mice, hM4Di n = 9 mice; interaction between hM4Di effect and null model effect, F_1, 15_ = 3.974, p = 0.0647. Sidak’s multiple comparison test: mCherry, Null versus Actual t_15_ = 6.026, ****p < 0.0001, hM4Di, Null versus Actual t_15_ = 3.485, **p = 0.0066. (l) Top: schematic diagram representing familiarity CCGP decoding performance between littermates and novel conspecifics. Bottom: both mCherry- and hM4Di-expressing mice showed significantly higher CCGP performances than CCGP performances from the null models. Two-way ANOVA with stacked matching: mCherry n = 8 mice, hM4Di n = 9 mice; interaction between hM4Di effect and null model effect, F_1, 15_ = 0.01545, p = 0.9027. Sidak’s multiple comparison test: mCherry, Null versus Actual t_15_ = 3.259, *p = 0.0105, hM4Di, Null versus Actual t_15_ = 3.638, **p = 0.0049. (m) Top: schematic diagram representing cross-familiarity CCGP decoding between familiar conspecific and littermate with novel conspecifics. Bottom: while mCherry-expressing mice showed significantly higher CCGP performances than CCGP performances from the null models, hM4Di-expressing mice showed comparable CCGP performances to the null models. Two-way ANOVA with stacked matching: mCherry n = 8 mice, hM4Di n = 9 mice; interaction between hM4Di effect and null model effect, F_1, 15_ = 4.858, *p = 0.0436. Sidak’s multiple comparison test: mCherry, Null versus Actual t_15_ = 2.565, *p = 0.0426, hM4Di, Null versus Actual t_15_ = 0.4923, p = 0.4923.

After confirming behavioral deficits by inactivating vCA1^→IL^ neurons in the extended social familiarization/recognition task, we analyzed social representations in IL^→NAcSh^ neurons. mCherry- and hM4Di-expressing mice exhibited comparable numbers of social cells (Supplementary Figs. 10e and 10f). Consistent with our results (Fig. 2i), a significantly higher overlap of social cells for familiar conspecifics and littermates (F∩L) was observed in vCA1 mCherry-expressing mice, but not in vCA1 hM4Di-expressing mice (Fig. 6e), suggesting that vCA1 inactivation induces alterations in neuronal representations of social targets in IL^→NAcSh^ neurons, which subsequently impairs social recognition. The LSVM classifier exhibited decent performances in predicting both social identity with an accuracy above the chance level even after inactivating vCA1^→IL^ neurons (Supplementary Fig. 10g).

Next, we examined the impact of vCA1 inactivation on the representation of social familiarity in IL^→NAcSh^ neural activities by using the CCGP classifier. The CCGP classifier was trained using data from interaction with familiar mouse 1 and novel mouse 2 on one side (e.g., F1^left^ and N2^left^) during the social recognition session and tested on data from familiar mouse 2 and novel mouse 1 on the opposite side (e.g., F2^right^ and N1^right^) (Fig. 6f). IL^→NAcSh^ neural activity showed high decoding performance for familiarity in mCherry-expressing mice even after CNO injection (Figs. 6g and 6i). However, consistent with behavioral data (Fig. 6d), vCA1 inactivation resulted in low CCGP performance for decoding familiarity using IL^→NAcSh^ neural activity in hM4Di-expressing mice (Figs. 6h and 6i). Moreover, CCGP classifier failed to decode social targets before and after familiarization after CA1 inactivation (F *vs.* F_N_, Fig. 6j). Both mCherry and hM4Di groups showed significantly high CCGP performances for classifying littermates from other social targets (L *vs.* F_N_, L *vs.* N; Figs. 6k and 6l). Lastly, to examine if the CCGP classifier could decode familiarity even when the familiarity levels between training and test sets differed, the classifier was trained using data from littermate and novel conspecific on one side (e.g., L1^left^ and N2^left^) and tested on data from familiar and another novel conspecific on the opposite side (e.g., F2^right^ and N1^right^). mCherry-expressing mice exhibited significantly higher decoding performance for familiarity, indicating that IL^→NAcSh^ neural activity represents social familiarity regardless of the familiarity level (Fig. 6m). However, in hM4Di-expressing mice, the CCGP classifier failed to distinguish between familiar and novel mice when the decoder was trained using data from littermates and novel mice (Fig. 6m). Meanwhile, position CCGP performances remained at the chance level for both mCherry- and hM4Di-expressing mice when comparing their neural representations between different social targets, suggesting that the neural representation from spatial information is not generalized in IL^→NAcSh^ neurons (Supplementary Figs. 11). Taken together, these results demonstrate that inactivating vCA1^→IL^ neurons after familiarization prevents the formation of representation of newly familiarized conspecifics without disrupting already stored social information in IL^→NAcSh^ neurons.

## Discussion

While the role of the hippocampus in social memory has been extensively investigated, it is not clear where social memory is stored in the neocortex. In this study, we found that a subpopulation of mPFC IL neurons stores consolidated memory of familiarized conspecifics including littermates and that the neural activity of vCA1^→IL^ neurons during offline phase is critical for these cortical social memory representations in male mice. Subsequently, these representations support the discrimination of familiar compared with novel conspecifics.

Previously, we have shown that IL^→NAcSh^ neurons are activated by familiar conspecifics and the activity of these neurons is required for social recognition, but not for object recognition in mice^25^. However, it remains unclear whether the neural activity of these neurons represents social memory, and if so, which aspects of social memory are encoded in these neurons and at which stage of social memory processing. We found that inactivating IL^→NAcSh^ neurons during the recognition of the social familiarization/recognition task, but not either the familiarization or consolidation offline phase, impairs social recognition 24 hours after familiarization, demonstrating that these neurons store long-term social memory and are required for the retrieval of social memory.

Our results go along with the complementary learning system model proposed by O’Reilly and colleagues, which suggests that hippocampal representations are gradually transformed into neocortical representations due to the temporal gap in learning between brain regions^3,31^. We found that inactivating vCA1 to IL neural projections during the offline consolidation phase impairs recognition memory of newly familiarized conspecifics without affecting memory of littermates, supporting the critical role of vCA1-IL interaction in consolidating newly established social memory. Previous studies have shown that dCA2 to vCA1 projections are crucial for encoding, consolidation, and recall of social memory^11^. Our study extends the previously known dCA2-vCA1 circuits for social memory by connecting the vCA1 to IL and uncovers differences in the temporal engagements and representational dynamics of social information in these brain regions. Our data further demonstrate the relatively slow learning within prefrontal subpopulations, which allows for the formation of neural representations of social targets 24 hours after familiarization. We also found that optogenetic inhibition of vCA1^→IL^ neurons during the social memory recognition session impaired recognition of newly familiarized conspecific. This suggests that the transfer of social memory from the hippocampus to the cortex may not be completed within 24 hours, as has been shown for other forms of memory^2^. Additionally, previous studies have shown that hippocampal dCA2 activity is necessary for recognizing littermates^10,11^, indicating that the hippocampus may play a sustained role in social memory retrieval. Whether the hippocampus eventually disengages at a remote time point or remains permanently engaged in social memory retrieval remains to be investigated.

We asked which aspects of social information are encoded by IL^→NAcSh^ neurons. We revealed that the activity of IL^→NAcSh^ neurons can simultaneously represent identity of familiar conspecifics as well as familiarity by turning the geometry of neural representations in neural activity space. Moreover, our results from TRAP experiments showing that inactivation of IL ^→NAcSh^ neurons responding to a familiar mouse impairs recognition of other familiar mice including littermates, strongly support that IL^→NAcSh^ neurons encode social familiarity. This aligns with previously established notion that learning induces generalization within prefrontal representations with potential mixed selectivity^6,32–35^.

The dCA2 was shown to be necessary for social memory during encoding, consolidation and retrieval phases^10,11^. In contrast to IL^→NAcSh^ neurons that are activated by familiar conspecifics, dCA2 neurons were shown to be activated by novel social targets^36^. However, a recent study showed that dCA2 neural activity can represent both littermates and novel mice in a higher- and lower-dimensional geometry, respectively^8^. The CCGP classifier successfully predict identities of novel mice, but CCGP for littermates was not greater than chance level, suggesting that CA2 neural activity supports generalized decoding of both identity and position only when subject mice explored novel mice^8^. Different from the dCA2, interestingly, we found that IL^→NAcSh^ neural activity supports greater generalization for the identity of littermates compared with novel mice. Also, CCGP classifier failed to decode the position of mice using IL^→NAcSh^ neural activities, regardless of their familiarity. These results show that the IL and dCA2 representations support the discrimination of novel versus familiar conspecifics in different ways. It is also worth noting that Hassan et al. recently showed that the generalization of the dCA2 neural representations is limited to distinguishing social and non-social odor cues^9^. Indeed, electrophysiological recordings in dorsolateral PFC and hippocampus of rhesus monkey revealed the potential capacities of generalization of contextual variables in both brain regions, eliciting the interplay of hippocampal-prefrontal cortex with their dynamical representations may be the key of social recognition^23^.

Curiosity still lies on the necessity of many brain regions and their connections in social memory processing. vCA1 sends its projections not only to the prefrontal cortex, but also to other brain regions including the NAc and amygdala via its collaterals^29,37^. Indeed, vCA1 to NAcSh projections have also been shown to store social memory in mice^14^. We speculate that distinct populations of vCA1 and their projections may either support either parallel processing of social information by sending different aspects of social information to each region or contribute to distributed representations of social information in multiple brain regions. For instance, a recent study reported that vCA1 neurons in mouse encodes identities, sex and strain of familiar conspecifics^38^. The hippocampus may serve as a critical hub for processing specific social information, which is then generalized and integrated within the mPFC through hippocampal-cortical interactions during consolidation.

Changes in neuronal ensembles in vCA1 supporting social memory have been reported in mouse models of autism spectrum disorder (ASD)^39,40^. Previously, we demonstrated that chronic social isolation during the juvenile period impairs social recognition in mice, which is accompanied by a selective reduction in the excitability of IL^→NAcSh^ neurons^25^. It would be valuable to examine the dynamics of neural representations of IL^→NAcSh^ neurons in ASD mouse models, as this could provide insights into the neural mechanisms underlying social recognition deficits in ASD.

## Methods

### Experimental model and subject details

Male C57BL/6NCrljOri mice were purchased from Orient Bio at PND 35 and introduced to the animal facility at Seoul National University College of Medicine. Four male mice were allocated into a standard mouse cage. Mice were habituated in the animal facility and may experience the surgical procedure at PND 42 depending on the experimental needs.

B6.Cg-Igs7^tm148.1(tetO–GCaMP6f,CAG–tTA2)Hze^/J(Ai148D) mice were obtained from the Jackson Laboratory (USA, Strain #:030328) and were bred in the specific-pathogen-free (SPF) animal facility at Seoul National University College of Medicine. Homozygous mutant males were mated with wild type female C57BL/6NSNU and their heterozygous offspring were weaned at PND 21 for the experiments. Four or five mice from the same parents were allocated into a standard mouse cage and raised until PND 42. Wildtype of Ai148 was used for miniscope recording with chemogenetic manipulation.

All mice were given by a fixed 12 hours light and dark cycle (light on: 8:00am – 8:00pm; light off: 8:00pm – next day 8:00am), while food and water were provided *ad libitum*. Experiments were conducted in accordance with the regulation of Seoul National University, College of Medicine, Department of Physiology and were approved by the Institutional Animal Care and Use Committee of Seoul National University (IACUC #: SNU-240103-10-1).

## Method details

### Surgical procedures

Subject mice were anesthetized with a Zoletil (30 mg/kg) and Rompun (10 mg/kg) mixture by intraperitoneal (i.p.) injection with the injection volume varied by subject body weight. Skin above mouse skull was gently rubbed with 70% (v/v) ethanol to sanitize while its fur on the corresponding area was shaved thoroughly before the anesthetized mouse was set on the rodent stereotaxic apparatus. Designated region of the mouse skull was drilled with a stereotaxically-fixed hand drill assembled with a 0.3 mm drill bit and virus injection was performed bilaterally.

The infralimbic cortex (AP +2.4 mm from bregma, ML ±1.0 mm from midline, DV -2.3 mm from skull; 10° angled injection to prevent virus expression in the prelimbic cortex), shell region of nucleus accumbens (AP +1.94 mm from bregma, ML ±0.55 mm from midline, DV - 4.15 mm from skull), and CA1 region of ventral hippocampus (AP -3.3 mm from bregma, ML ±3.1 mm from midline, DV -3.9 mm from skull) were targeted for virus injection. For the virus injection, a glass capillary (#504949, WPI, USA) was pulled out with appropriate diameter using the Dual-Stage Glass Micropipette Puller (NARISHIGE, USA) and filled with mineral oil (M5904, Sigma-Aldrich, USA), while virus was filled from the front side of the glass capillary. All viruses were purchased from Addgene (USA) except AAV_8_-fos^CreERT^^2^-PEST (Stanford GVVC, USA).

For GCaMP6f expression in Ai148 mouse, AAV_rg_-Ef1a-mCherry-IRES-Cre was injected as 5×10^12^ GC/mL (250 nL) into the NAcSh. For TRAP labeling, AAV_8_-fosCreER^T2^-PEST was injected as 1×10^13^ GC/mL (200 nL) in IL and AAV_rg_-DIO-hM4Di-mCherry or AAV_rg_-DIO-mCherry (for control) was injected as 5×10^12^ GC/mL (200 nL) in NAcSh. For optogenetics, AAV_5_-hsyn-NpHR3.0-YFP or AAV_5_-hsyn-eYFP (for control) was injected in IL or vCA1 as 2.5×10^12^ GC/mL (200 nL). For chemogenetic manipulation, AAV_PHP.eB_-hsyn-DIO-hM4Di-mCherry or AAV_8_-hsyn-DIO-mCherry (for control) as 5×10^12^ GC/mL (200 nL) was injected in IL or vCA1, while AAV_rg_-Ef1a-mCherry-IRES-Cre was injected as 5×10^12^ GC/mL (200 nL) into NAcSh or IL. For the tracing, AAV_8_-DIO-synaptophysin-mRuby as 5×10^12^ GC/mL (200 nL) was injected in vCA1 while AAV_rg_-hsyn-HI-eGFP-Cre with same titer and volume was injected in either IL or NAcSh. For the miniscope recording with vCA1 chemogenetic inactivation, wildtype mice from Ai148 mouse line were used and injected with AAV_rg_-Ef1a-Flpo, AAV_2_-flex-GCaMP6f mixed with 1:1 ratio with each titer 5×10^12^ GC/mL into IL (250 nL). AAV_rg_-Ef1a-Cre was injected as 5×10^12^ GC/mL (200 nL) into NAcSh and AAV_8_-hsyn-fDIO-hM4Di-mCherry or AAV_8_-hsyn-fDIO-mCherry (for control) was injected as 5×10^12^ GC/mL (300nL) in vCA1. The virus was diluted with filtered ACSF to the desired titer before use. The virus was delivered to the target region of interest with a rate of 20-30 nL/minute. After the virus injection, glass capillary was held still within the brain at least 5 minutes before being slowly withdrawn. The incised skin was sutured with sterilized suture thread and applied with additional Betadine to prevent any infection. Mice were placed on a heating pad until the anesthesia wore off and were then returned to their home cages for further rest.

### Integrated GRIN lens preparation

To observe neural activity from the pyramidal layer of IL cortex while preventing any damage to the prelimbic cortex, a GRIN lens with a 1 mm diameter and 4 mm length (Inscopix, USA) was attached to a 1 mm right-angle prism with an aluminum coat (#86-621, Edmund optics, USA) using a droplet of optical adhesive (NOA65, Edmund optics, USA). The assembly was UV-cured for at least 10 minutes and then immersed in 70% ethanol to sterilize before use.

### Ca^2+^ imaging surgery

Two weeks after the virus injection, the mice underwent an additional stereotaxic surgery for Ca2+ imaging. First, a sterilized steel screw with a 3 mm diameter was fastened to the skull before performing the craniotomy. To implant the GRIN lens, a 1.3 mm × 1.3 mm rectangular craniotomy was made above the IL viral injection site. The cortical tissue adjacent to the IL was carefully aspirated using 28-gauge blunt needles. Cold saline was constantly applied during the cortex aspiration to prevent tissue desiccation and further cell death. After aspirating 1.5 mm of cortical tissue, a custom-designed 1.2 mm laser-cut disposable blade was inserted into the cortex, adjacent to the IL, to create an additional 1 mm sheath for the integrated GRIN lens. After inserting the laser blade and gently moving it away from the midline, cold saline was applied to irrigate any excess bleeding. Once bleeding was confirmed to have stopped, the integrated GRIN lens was quickly inserted into the site to a depth of 2.5 mm. Following lens implantation, tissue adhesive (3M Vetbond) was applied to the junction between the GRIN lens and the skull, and dental cement (SUN MEDICAL, Japan) was applied in an appropriate amount to secure the lens in place and cover the exposed skull. After allowing at least 15 minutes for the dental cement to harden, a baseplate for miniscope recording was placed on top of the GRIN lens and cemented into position. A plastic dummy cap was then locked onto the baseplate to prevent photobleaching and lens damage. Dexamethasone (0.3 mg/kg) was administered during surgery and continued for a week post-surgery.

### Drug cannula implantation

To induce local inactivation of neural terminals using chemogenetics or optogenetics, a drug cannula or optic cannula was implanted at the target site. Two weeks after the virus injection, the mice underwent an additional stereotaxic surgery for cannula implantation. To begin, a sterilized steel screw with a 3 mm diameter was fastened to the skull to prevent any artifact from the movement of subject. The designated region of the mouse skull was drilled bilaterally using a stereotaxically-fixed hand drill with a 0.5 mm drill bit. A guide cannula (#62003, RWD, USA) for micro-infusion and an optic cannula (#R-FOC-L200C-39NA, RWD, USA) for local optogenetic silencing were slowly implanted at the target site using a 3D-printed cannula holder. After the implantation, tissue adhesive was applied to the gap between the bottom of the cannula and the attached skull, ensuring it did not spread to other skull holes. After waiting for 5 minutes to allow the adhesive to fully dry, the cannula was gently detached from the cannula holder. The other cannula was implanted in the remaining hole with a similar slow approach, and tissue adhesive was applied before the cannula holder was removed.

After the cannula implantation, dental cement was applied in an appropriate amount to secure the cannula in the desired location and to cover the exposed skull. Once the dental cement had fully dried, a sterilized plastic dummy cap (#62102, RWD, USA) was inserted and tightly joined with each guide cannula to prevent any clogging of the guide cannula pathway during recovery. Dexamethasone (0.3 mg/kg) was administered during surgery and for three consecutive days post-surgery. During recovery, the mice may nibble on and remove the dummy cap, so constant monitoring was necessary. If the dummy cap was damaged or removed, it was replaced with a new sterilized dummy cap, which was tightly secured to the guide cannula.

### CNO administration

Clozapine N-oxide (CNO) dihydrochloride (#HB6149, Hellobio, UK) was used to activate designer receptors exclusively activated by designer drugs (DREADDs). CNO was dissolved in 0.9% normal saline to yield a stock concentration of 4 mg/mL and stored in a -20 ℃ freezer for up to 6 months. For i.p. injection, CNO was diluted with 0.9% normal saline to achieve a final concentration of 1 mg/mL and injected into subject mice to obtain a 3 mg/kg body concentration. The same volume of saline was injected for control. Behavioral assessments were started 40 minutes after drug injection.

For local administration, the subject mouse was isolated into a cage with clean bedding, and 500 nL of 1 mM diluted CNO solution was infused per site over 5 minutes using a syringe pump (Pump11, Harvard Apparatus, USA) mounted with a 10 µL syringe (#701, Hamilton, USA), allowing the mouse to move freely during the infusion. The infusion cannula (#62203, RWD, USA) was gently removed, and the original dummy cap was tightly secured 3 minutes after the end of the micro-infusion to prevent drug reflux. The subject mouse was then returned to its home cage. To prevent any side effects from drug hydrolysis, at least a two-day interval was given between treatments with counterbalanced drugs.

### Social target – specific engram labelling using TRAP

We employed viral-mediated targeted recombination in active population (TRAP) technology to specifically tag neurons activated during interactions with a particular social target. Experimental procedures from previous study were carried out^27^. On the day of experiment, 4-hydroxytamoxifen (4-OHT; #H6278, Sigma-Aldrich, USA) was dissolved at 2 mg/ml with 2 % tween-80 (#P1754, Sigma-Aldrich, USA) and 5 % DMSO (#D2650, Sigma-Aldrich, USA) in 0.9 % normal saline. For TRAP tagging, subject mice were habituated to home cage isolation for 1 hour each day for 5 days. On day 6, each subject mouse was isolated, and a perforated acrylic box containing a novel social target was introduced to allow the subject to investigate the novel conspecific for 30 minutes. After 30 minutes, the target mice were returned to their home cages, while the subject mice remained isolated for an additional 30 minutes for a post-learning session before being returned to their home cages.

On the day of the experiment, the subject was isolated 30 minutes prior to social investigation. As on the previous day, the same social target was introduced, this time without a perforated chamber, for 10 minutes. During this 10-minute period, subject mice that exhibited excessive aggression towards the social target were excluded from further experiments. After 10 minutes, the target conspecific was removed from the cage, while the subject mouse remained in the cage for an additional hour. Following this, the subject mouse was injected with the prepared 4-OHT (body concentration: 10 mg/kg) via i.p. injection to tag neurons that had been activated during the social interaction. Two hours after the 4-OHT injection, the subject mice were returned to their home cages. To induce the expression of the fos-dependent hM4Di designer protein, behavior assessments were started one week after the TRAP tagging procedure.

### Behavior Tests

All behavior tests described below were performed in a soundproof chamber with a dim light illumination within mouse light-on cycle. All behavior tests were recorded by the camcorder while the experimenters were blinded to the experimental conditions. Both the subject and target mice were habituated with handling prior to any behavior test. For the optogenetic inactivation of IL^→NAcSh^ neurons or vCA1^→IL^ neurons, the bilaterally implanted optic fibers located above either the NAcSh or IL were connected to a laser (594nm wavelength at 6mW), with mating sleeves. The same behavior paradigm was repeated to assess the effect of inactivating the target circuit during whole or part of behavior paradigm, as well as the effect of given laser treatment. The laser was administered to the subject mice in a randomly counterbalanced manner. For social familiarization/recognition task, same batch of mice were used to observe the effect of neural inactivation during social familiarization or recognition session, except an additional batch for vCA1^→IL^ inactivation during social recognition (Fig. 5e and Supplementary Fig. 7c). For the necessary Ca^2+^ imaging, each subject mouse was fully habituated with handling and miniscope application for at least a week prior to the Ca^2+^ imaging. A custom labview code was used to synchronize mice behavior monitored by Basler camera and Ca^2+^ activity while active commutator being performed to prevent any tension on the coaxial cable of miniscope. Mice behavior was manually analyzed by an experienced blind experimenter.

### Social familiarization/recognition task

Social familiarization/recognition task took 2 days to observe the functional role of specific neural circuit at different stages of memory consolidation. Prior to the behavior task, subject mouse was isolated in a cage with fresh bedding. Subject mice were placed in a white opaque linear acrylic box (10 × 60 cm size box)^41^ and allowed freely moving exploration. After 5 minutes of familiarization, a target mouse placed in a perforated acrylic chamber was introduced in either left or right side randomly to prevent location bias. After 5 minutes of interaction, subject and target mice were returned to each isolation cage for a rest. After 5 minutes of rest, subject mouse was reallocated to the linear chamber with same target mouse in the perforated chamber in different side for second interaction and repeated once more. After social interaction, both subject and target mouse returned to their original cage. After 24 hours, subject mice underwent social recognition session in the same linear chamber for the familiarization. Subject mice explore a perforated chamber with familiar, novel conspecific or littermate with random order and 5 minutes of interval between each interaction session. Depend on the need, subject mouse might be attached with the integrated GRIN lens and monitor the activity of IL^→NAcSh^ neuron with open-source miniscope (UCLA miniscope V4, Open ephys, USA) and CNO may be injected by I.P.

### Consecutive littermate recognition task

Same behavior chamber from the social familiarization/recognition task was used to carry the consecutive littermate recognition task. At least two days prior to the behavior tests, littermates that were planned as target mice were habituated to the interaction chamber and the subject mouse was allowed to watch its littermates located in the perforated chamber. This prevents littermates from expressing unwilling empathy towards their littermates so that the littermates exhibiting unusually high interaction times towards their known littermates. On the test day, subject mouse was isolated in a cage with fresh bedding prior to the behavior task. Subject was placed in the behavior chamber and allowed freely moving exploration. During familiarization session, two of subject’s littermate were isolated as target mice in separate cages with fresh bedding prior to the behavior task. After 5 minutes of familiarization, one of the target mice was placed in the perforated acrylic chamber in either left or right side. After 5 minutes of interaction, subject and target mice were returned to each isolation cage to take a rest. Then, subject mouse was reallocated to the linear chamber with an alternative target mouse in the perforated chamber in different side for second interaction. Subject interacting with its littermates was repeated once more (two more littermate interaction) with different position of linear track compared to the initial session. After social interaction, both subject and target mouse returned to their original cage.

### Consecutive novel mouse interaction task

Consecutive novel mouse interaction task was carried similar to the consecutive littermate recognition task, but with two novel conspecifics as target mice.

### Open field test

A subject mouse was placed in the center of a white opaque acrylic box (20 cm diameter cylinder) to move freely for 5 minutes, consecutively followed by an additional 5 minutes of yellow-light session. The order of yellow-light session and no-light session were randomly counterbalanced.

### Real time place preference assay

A subject mouse was placed in the center of a white opaque acrylic box (33×33 cm) to move freely for 10 minutes. During exploration, the location of subject mouse was lively tracked.

When the subject mouse entered a randomly set quadrant of the behavior chamber, live monitoring system sent an external TTL signal towards 594nm yellow laser (ADR-800A, Shanghai Laser and Optics Century, China) so that each cannula was intended to receive 10 mW of 594nm yellow light if the subject mouse entered an intended quadrant area.

### Novel object recognition test

Training and test session of novel object recognition test were conducted at the same day. A subject mouse was placed in the center of a white opaque acrylic box (33×33 cm) to move freely for 10 minutes for familiarization. After the familiarization session, two identical objects were placed on the behavior chamber so that the subject mouse can investigate two novel objects for 10 minutes training session. After the training session, the subject mouse was taken out from the behavior chamber as one of the objects was randomly replaced with a novel object. To prevent from providing any scent cue to the subject mouse, the original object which was placed inside the test area was wiped with 70% ethanol. Right after the object replacement, the subject mouse was reintroduced into the behavior chamber to explore for 10 minutes. To investigate the effect of chemogenetic inactivation, each subject mouse underwent novel object recognition test twice with two different pairs of objects.

### Immunohistochemistry

After experiment, each brain of subject mouse was extracted after transcardial perfusion by cold PBS solution followed by 4% PFA solution (T&I, Korea). Perfused brains were emerged into 4% PFA and stored in 4 °C for 24 hours. Then, the fixed brains were emerged again into 30% (w/v) sucrose solution for 48 hours in 4 °C. Processed brain was frozen at -20 °C with frozen section compound (#FSC22, Leica, Germany) and its 30 μm thickness brain slices were stored in CPS solution for the next usage. To stain cFos, brain slices were incubated in a blocking solution with 4% (v/v) goat serum, 0.1% (v/v) Triton X-100 and PBS for 40 minutes. Brain slices were then incubated with primary antibody (#2250, Cell signaling Technology, USA) above in 4°C 30 RPM locker for the next 48 hours. After that, brain slices were incubated with Alexa Fluor 488 donkey anti-Rabbit IgG secondary antibody (#A-21206, Invitrogen, USA) on room temperature for 4 hours and washed with PBS. 0.01% (v/v) of DAPI (#62247, Thermo Scientific, USA) was used for counterstaining. Fluorescent image was obtained using Axio Scan Z1 (Zeiss, Germany) and the fluorescent labeled cells were counted manually after blind the identity of mouse. Ca^2+^ data processing

After obtaining 20-30 FPS Ca^2+^ imaging video from the open-source miniscope during rodent behavior, videos were concatenated and converted to tiff format with software Image J. Custom-modified CNMF-e was used to extract fluorescence signal from individual regions of interests (ROIs). For each trial from a subject, Ca^2+^ transients from custom-modified CNMF-e was used to analyze neuronal activity with further process. cell activities from multiple days were registered showing more than 80% of preserved region of interest (ROI) without discernible GCaMP expression artifacts (Supplementary Figs. 2c-f, 5a-5d, 10a-10d). Ca^2+^ transients from each subject were synchronized with the behavior data obtained with manual scoring. Behavior data was classified with binary label with either 0 or 1, as 0 means no interaction and 1 means the subject being under social interaction. 11 Ai148 mice in Fig. 2 exhibited 558 social cells with the total 1181 detected cells. 8 mCherry - expressing mice in Fig. 6 exhibited 454 social cells with the total 1038 detected cells, while 9 hM4Di - expressing mice exhibited 361 social cells with the total 862 detected cells.

### Receiver operating characteristic (ROC) analysis

To identify whether individual cell from IL^→NAcSh^ neuronal population is reactive to social behavior, all Ca^2+^ traces were z-scored and a receiver operating characteristic (ROC) analysis for each neuronal activity based on the range of binary thresholds of social stimulus was calculated^17^. ROC analysis was followed to identify binary event vector based on mice behavior to calculate the true positive rate (TRP) and the false positive rate (FPR) of individual neurons. ROC curve was drawn based on the TRP against the FPR over the binary threshold range^17^. The area under the ROC curve (auROC) of a neuron was compared to the 1,000 auROC obtained from randomly circular permuted Ca^2+^ signals of the same cell. A neuron was considered to be significantly responsive if its auROC value is over the 95^th^ percentile of the random distribution, and labeled as “social cells” in this research. The identities of social cells were recorded and compared with different interaction sessions to observe the overlapping social cell population within a behavioral paradigm.

To examine the effect of familiarization, the interaction-dependent Ca^2+^ AUCs of social cells from familiar mouse were obtained during social familiarization (F_N_, F_N_’, and F_N_’’) and recognition (F). To determine whether the reactivity of social cells depends on familiarity, the interaction-dependent Ca^2+^ AUCs were calculated individually for social cells from familiar conspecifics (F), novel conspecifics (N), and littermate (L) during interactions with corresponding social targets.

### Population decoding analysis

The decoding analysis for the population activities was performed with reference to what was described^8,42^. A linear classifier based on a support vector machine (SVM) was used with custom-written matlab and python scripts based on the scikit-learn SVC package for the cluster visualization^43^.

To make pseudo-trials, Ca^2+^ transient data from total neuron population which correspond to social interaction bouts were used and divided into 200 ms time bins. Mean event activity was calculated by averaging all Ca^2+^ transients within each bin. As 75% of pseudo-trial data was used to train the classifier, remaining 25% of data was used to test the trained classifier to get decoding performance within subject. Pseudo-population activity vectors were obtained from both the training and testing datasets. Randomly sampled *q* population vectors (where *q* = 5 otherwise noted) from all n neurons in the training dataset for each subject were used to concatenate them into a single *qn*-long vector, where *n* is the total number of recorded neurons in each subject. Same procedures were repeated to construct a training dataset of pseudo-population activity vectors. For a meaningful comparison of decoding results across experiments, only subjects that explored each session for more than 3 seconds each were included in the decoding analysis^8^.

To assess the statistical significance of the decoding performance, a null model in which the binary labels (0 or 1 as described above) of pseudo-trials were randomly shuffled. Same procedures were repeated for each decoder to obtain the decoding performance with null-model.

### Dimensional reduction and cluster metrics

To visualize the pseudo-population activity vectors on a reduced-dimensional space (*dimension* = 3 unless specified), a principal component analysis (PCA) was performed on the training and testing datasets and the result of dimensional reduction was projected on a space.

For each trial, the mean of each cluster sample (*x*) as the position of the corresponding centroid (*C*_*t*_) was defined as the following expression:

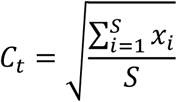

where *t* was trial and *S* was the number of samples in the reduced dataset.

The mean distance between all samples to the centroid for each cluster as the radius (*R*_*t*_) of the corresponding cluster was calculated by the following expression:

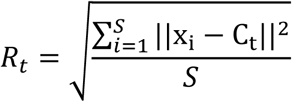

where the ||x_i_ − C_t_|| represented the Euclidean distance from the centroid to samples.

### Cross-condition generalization performance (CCGP)

CCGP was computed as described^8,42^. To make training and testing datasets, pseudo-population activity vectors for each trial was constructed to extract features as described above. To carry CCGP, extracted features from experimental datasets were grouped into two pairs – training and testing datasets. Each training and testing dataset contained binary labels (0 or 1) corresponding to the decoding trial conditions (identity or position). CCGP was calculated by averaging the decoding performances from *e* epochs (where *e* = 5 unless noted) with randomly extracted two pairs of training and testing datasets. For CCGP analysis, the training and testing datasets were reversed, and the decoding performance was averaged, except the CCGP analysis for cross-position. For example, identity CCGP for littermates was measured by training a linear classifier to decode the identity of two littermates on the left side (L1^left^ and L2^left^) and tested on data from littermates on right side (L1^right^ and L2^right^). The process was repeated by training the classifier with data from the littermates on the right side and testing it on data from the left side, with the decoding performances averaged together. CCGP analysis for cross-familiarity was done based on the target conspecific’s orientation. For instance, CCGP was measured by training a linear classifier to decode the identity of two conspecifics on the left side (L1^left^ and N2^left^) and testing it on data from other conspecifics on right side (F2^right^ and N1^right^). Additionally, the decoding performance of the linear classifier was tested by training it on two conspecifics on the right side (L2^right^ and N1^right^) and testing it on data from left side (F1^left^ and N2^left^). The two CCGP performances were averaged. For the cross-position analysis, CCGP was measured by training a linear classifier to decode the position of two known conspecifics (F1^left^ and L2^right^) and tested on data from two novel conspecifics _(N1left and N2right)_

### Quantification and statistical analysis

Paired t-tests, one-sample t-test, one-way and two-way repeated-measure analysis of variance (ANOVA) were followed by appropriate multiple comparisons post hoc to analyze mouse behavior and Ca^2+^ data.

### Data availability

All raw data and reagents are available from the corresponding author upon reasonable request.

### Code availability

All software or algorithm used in this study is available and described in the method section. The custom-written Python scripts used in this manuscript is available at the GitHub repository (https://github.com/minsmis/Social_familarity.git).

## Acknowledgements

This work was supported by grants to Y.-S.L (NRF-2023R1A2C200322912 and NRF-2022ME5E801804913), S.J.K. (NRF-2018R1A5A202596423), and G.P (RS-2023-00271950) from the National Research Foundation of Korea. The authors thank Dr. Sung-Phil Kim, Dr. Paul Frankland, and all the lab members for their valuable comments on the manuscript, and Taewoo Kim, Min-Gyun Kim, and Yeji Song for assisting rodent stereotaxic surgeries and behavior analysis.

## Author contributions

Y.-S.L. and G.P. conceived the study and designed the experiments with contribution from S.J.K. and D.L. G.P. performed most of the experiments. G.P. performed most of the data analyses with advice from M.S.K., Y.-B.L., D.L and Y.-S.L. S.S. wrote codes for miniscope data acquisition. G.P. and Y.-S.L. wrote the manuscript with input from M.S.K. Y.-B.L. S.J.K. and D.L.

