## Supplementary figures and legends for "Hippocampal-cortical interactions in the consolidation of social memory"

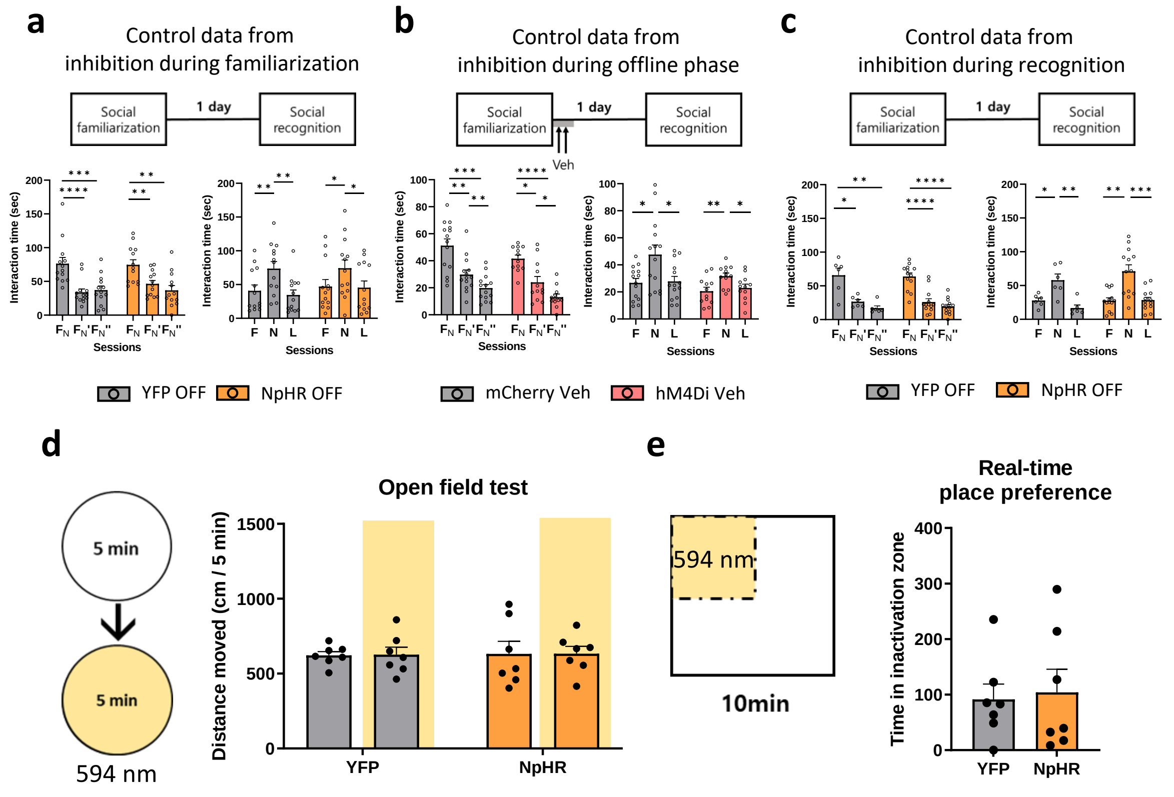


### Supplementary Figure 1.

**Supplementary Figure 1. Control behaviors with or without inactivation of IL$^{\rightarrow NAcSh}$ neurons**

(a) Social familiarization without optogenetic manipulation. Left: both YFP- and NpHR-expressing mice showed significantly decreased interaction times during the social familiarization session when the laser inactivation was not applied. One-way ANOVA with repeated measures: YFP, n = 13 mice; $F_{1.475, 17.7} = 35.11$, ****$p < 0.0001$. Tukey's multiple comparison test: $F_N$ versus $F_N$' $q_{12} = 10.27$, ****$p < 0.0001$; $F_N$ versus $F_N$'' $q_{12} = 8.089$, ***$p = 0.0003$; $F_N$' versus $F_N$'' $q_{12} = 1.230$, $p = 0.6685$. NpHR, n = 13 mice; $F_{1.486, 17.83} = 11.54$, **$p = 0.0013$. Tukey's multiple comparison test: $F_N$ versus $F_N$' $q_{12} = 6.720$, **$p = 0.0013$; $F_N$ versus $F_N$'' $q_{12} = 5.270$, **$p = 0.0075$; $F_N$' versus $F_N$'' $q_{12} = 1.722$, $p = 0.4658$. Right: YFP- and NpHR-expressing mice showed significantly longer interaction times with novel conspecifics during the social recognition when laser inactivation was not applied in the social familiarization session. One-way ANOVA with repeated measures: YFP, n = 13 mice; $F_{1.606, 19.28} = 17.12$, ***$p = 0.0001$. Tukey's multiple comparison test: F versus N $q_{12} = 6.814$, **$p = 0.0011$; F versus L $q_{12} = 1.553$, $p = 0.5332$; N versus L $q_{12} = 6.355$, **$p = 0.0020$, NpHR, n = 13 mice; $F_{1.748, 20.97} = 6.936$, **$p = 0.0063$. Tukey's multiple comparison test: F versus N $q_{12}$ = 4.093, *p = 0.0335; F versus L $q_{12}$ = 0.3644, p = 0.9642; N versus L $q_{12}$ = 4.259, *p = 0.0271.

(b) Social memory consolidation without chemogenetic manipulation. Left: both mCherry- and hM4Di-expressing mice showed significantly decreased interaction times during the social familiarization session. One-way ANOVA with repeated measures: mCherry, n = 14 mice; $F_{1.348, 17.52}$ = 23.54, ****p < 0.0001. Tukey's multiple comparison test: $F_N$ versus $F_N$' $q_{13}$ = 5.588, **p = 0.0044; $F_N$ versus $F_N$'' $q_{13}$ = 8.165, ***p = 0.0002; $F_N$' versus $F_N$'' $q_{13}$ = 5.473, **p = 0.0051, hM4Di, n = 12 mice; $F_{1.201, 13.21}$ = 20.40, ***p = 0.0004. Tukey's multiple comparison test: $F_N$ versus $F_N$' $q_{11}$ = 4.104, *p = 0.0354; $F_N$ versus $F_N$'' $q_{11}$ = 13.30, ****p < 0.0001; $F_N$' versus $F_N$'' $q_{11}$ = 3.987, *p = 0.0408. Right: both mCherry and hM4Di mice showed distinct preference towards novel conspecifics during the social recognition task when injected with vehicle in the offline phase. One-way ANOVA with repeated measures: mCherry, n = 14 mice; $F_{1.369, 17.80}$ = 8.918, **p = 0.0046. Tukey's multiple comparison test: F versus N $q_{13}$ = 4.770, *p = 0.0129; F versus L $q_{13}$ = 0.5037, p = 0.9328; N versus L $q_{13}$ = 4.199, *p = 0.0274, hM4Di, n = 12 mice; $F_{1.748, 19.23}$ = 7.769, **p = 0.0044. Tukey's multiple comparison test: F versus N $q_{11}$ = 5.957, **p = 0.0038; F versus L $q_{11}$ = 0.9779, p = 0.7732; N versus L $q_{11}$ = 4.556, *p = 0.0205.

(c) Social recognition without optogenetic manipulation. Left: both YFP and NpHR mice showed significantly decreased interaction times during the social familiarization session. One-way ANOVA with repeated measures: YFP, n = 6 mice; $F_{1.374, 6.868}$ = 16.28, **p = 0.0037. Tukey's multiple comparison test: $F_N$ versus $F_N$' $q_5$ = 4.917, *p = 0.0395; $F_N$ versus $F_N$'' $q_5$ = 7.414, **p = 0.0077; $F_N$' versus $F_N$'' $q_5$ = 2.357, p = 0.3042. NpHR, n = 13 mice; $F_{1.958, 23.49}$ = 44.28, ****p < 0.0001. Tukey's multiple comparison test: $F_N$ versus $F_N$' $q_{12}$ = 9.947, ****p < 0.0001; $F_N$ versus $F_N$'' $q_{12}$ = 12.18, ****p < 0.0001; $F_N$' versus $F_N$'' $q_{12}$ = 1.993, p = 0.3672. Right: both YFP and NpHR mice showed significantly longer interaction times with novel conspecifics during the social recognition session when laser inactivation was not applied in the social recognition session. One-way ANOVA with repeated measures: YFP, n = 6 mice; $F_{1.186, 5.928}$ = 20.91, **p = 0.0033. Tukey's multiple comparison test: F versus N $q_5$ = 5.230, *p = 0.0315; F versus L $q_5$ = 5.590, *p = 0.0244; N versus L $q_5$ = 7.768, **p = 0.0063, NpHR, n = 13 mice; $F_{1.201, 14.41}$ = 24.13, ***p = 0.0001. Tukey's multiple comparison test: F versus N $q_{12}$ = 6.854, **p = 0.0011; F versus L $q_{12}$ = 0.2472, p = 0.9833; N versus L $q_{12}$ = 7.615, ***p = 0.0004.

(d) Effect of inactivation of $IL^{\rightarrow NAcSh}$ neurons on open field test. Left: schematic diagram of open field test (OFT) with NpHR inactivation. Yellow light (594 nm, 6 mW) was illuminated bilaterally for 5 minutes with counterbalancing. Right: Inactivation of IL$^{\rightarrow NAcSh}$ neurons did not affect mice basal locomotor activity. Ordinary two-way ANOVA: YFP, n = 7 mice; NpHR, n = 7 mice, $F_{1, 24}$ = 0.001579, p = 0.9686. Sidak's multiple comparisons test: YFP, x light – light $q_{24}$ = 0.07069, p = 0.9969; NpHR, x light – light $q_{24}$ = 0.01449, p = 0.9999

(e) Effect of inactivation of IL$^{\rightarrow NAcSh}$ neurons on RTPP test. Left: schematic diagram of real-time place preference (RTPP) with IL$^{\rightarrow NAcSh}$ inactivation. Right: inactivation of IL$^{\rightarrow NAcSh}$ neurons did not affect mice place preference. Unpaired t-test: YFP, n = 7 mice; NpHR, n = 7 mice, $t_{12}$ = 0.2572, p = 0.8014.

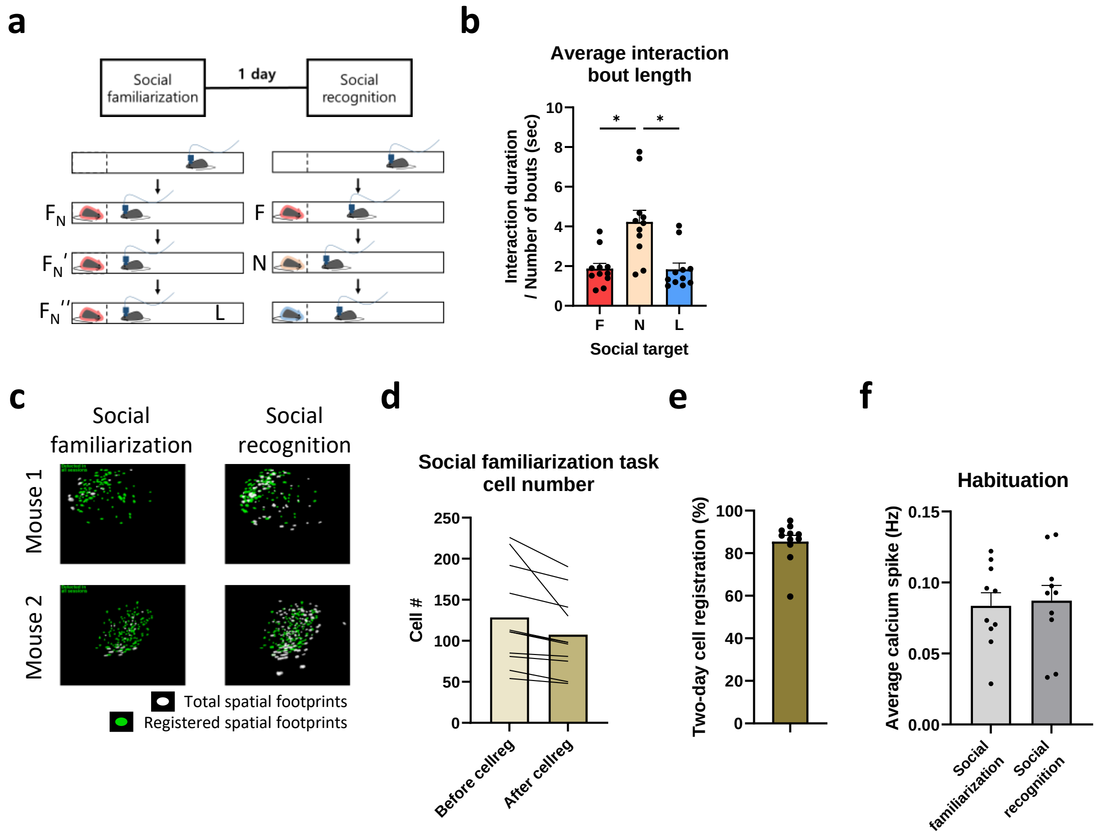


**Supplementary Figure 2.**

**Supplementary Figure 2. Imaging *in vivo* $Ca^{2+}$ activity of $IL^{\rightarrow NAcSh}$ neurons during the social familiarization/recognition task**

(a) Experimental scheme for the social familiarization/social recognition task with miniscope imaging. After habituation (5 min), a subject mouse was familiarized to a conspecific ($F_N$) three times for 5 minutes, with two 5-minute intervals during the social familiarization session. On the next day of social recognition session, the subject mouse interacted with familiarized social target (F), a novel conspecific (N), or a littermate (L).

(b) Average social interaction bout lengths of subject mice were different depending on the familiarity of social targets. Average interaction bout length was calculated by dividing total interaction duration with interaction frequency. One-way ANOVA with repeated measures, $F_{1.435, 14.35} = 8.910$, $^{**}p = 0.0055$; Tukey's multiple comparison test: F versus N $t_{10} = 4.422$, $^{*}p = 0.0265$, F versus L $t_{10} = 0.1154$, $p = 0.9963$, N versus L $t_{10} = 4.604$, $^{*}p = 0.0215$.

(c) Representative images from two subject mice showing longitudinal cell registrations from ROIs over multiple days. Among the total spatial footprints (white), footprints marked with green color represent registered cells.

(d) Longitudinal cell registration resulted in identification of smaller but significantly reduced cell populations within social familiarization session. Paired t-test, $t_{12} = 2.210$, *$p = 0.0423$.

(e) 85.45% of neuronal populations were successfully registered across multiple days.

(f) $Ca^{2+}$ transients during each habituation session were comparable between the social habituation and recognition session. One-way ANOVA with repeated measures: n = 11 mice; $F_{2.587,25.87} = 0.6541$, $p = 0.5661$.

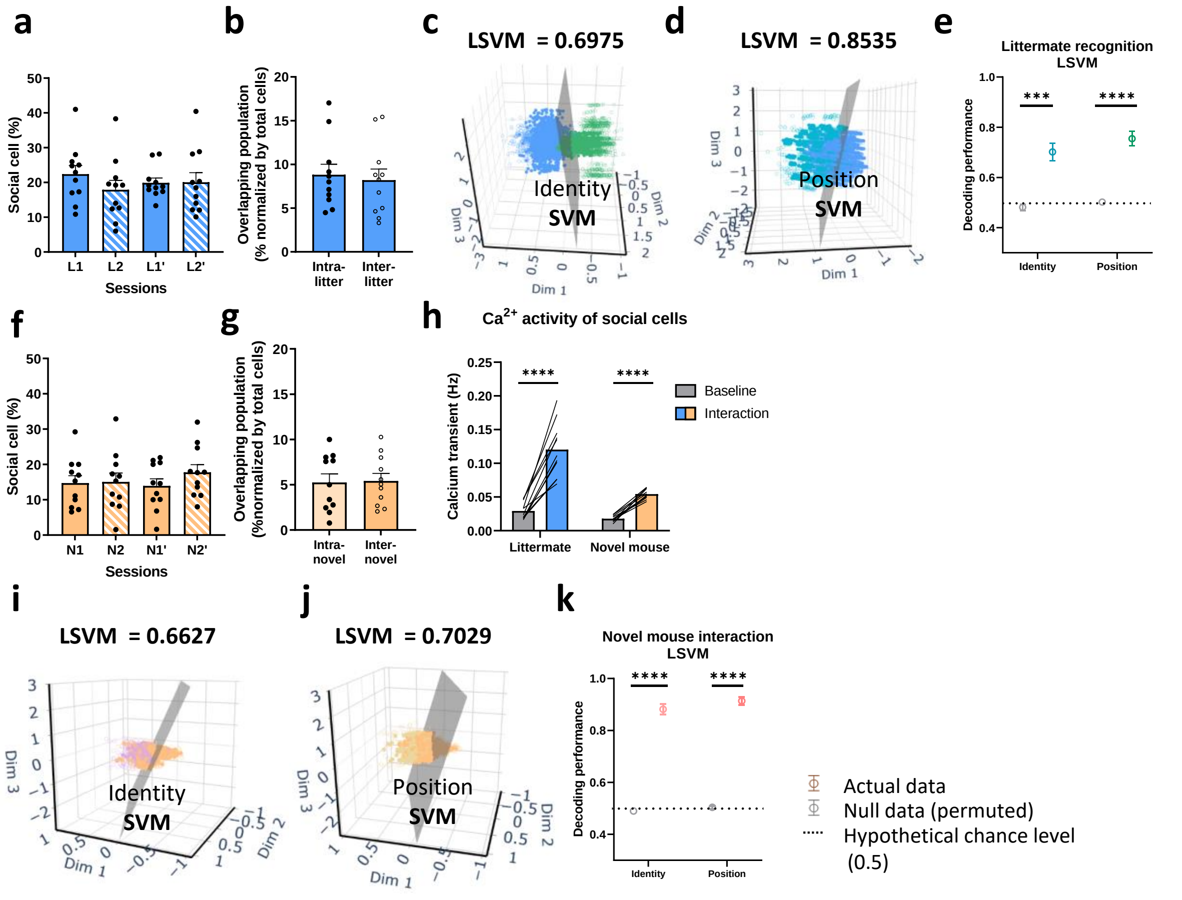


### Supplementary Figure 3.

**Supplementary Figure 3. Overlapping social cell analysis, identity and position decoding using LSVM classifier**

(a) The numbers of social cells from the littermate recognition task are comparable among sessions. One-way ANOVA with repeated measures: n = 11 mice; $F_{1.870, 18.70} = 0.6805$, p = 0.5090.

(b) Overlapping populations of social cells from different littermates (Inter-litter) or the same littermate (Intra-litter) were comparable in $IL^{\rightarrow NAcSh}$ neurons. Two-tailed paired t-test: n = 11 mice; $t_{10} = 0.9040$, p = 0.3873.

(c) A representative example of visualizing the 3D-LSVM where the activities of $IL^{\rightarrow NAcSh}$ neurons decode the identity of littermate.

(d) A representative example of visualizing the 3D-LSVM where the activities of $IL^{\rightarrow NAcSh}$ neurons decode the position of littermate.

(e) Both identity and position information of littermate were well classified using the 3D-linear SVM model compared to those from null models. Identity, Two-tailed paired t-test: n = 11 mice; $t_{10}$ = 5.293, ***p = 0.0004. Position, Two-tailed paired t-test: n = 11 mice; $t_{10}$ = 7.989, ****p < 0.0001.

(f) The numbers of social cells from novel conspecific interactions are comparable among sessions. One-way ANOVA with repeated measures: n = 11 mice; $F_{1.935,19.35}$ = 0.5940, p = 0.5566.

(g) Overlapping populations of social cells from different littermates (Inter-novel) or the same littermate (Intra-novel) were comparable in $IL^{\rightarrow NAcSh}$ neurons. Two-tailed paired t-test: n = 11 mice; $t_{10}$ = 0.3684, p = 0.7203.

(h) Social cells from littermate interactions showed significantly increased $Ca^{2+}$ activity during social interaction (left). Two-tailed paired t-test: n = 11 mice; $t_{10}$ = 8.693, ****p < 0.0001. Social cells from novel mouse interactions also showed significantly increased $Ca^{2+}$ activity during social interaction (right). Two-tailed paired t-test: n = 11 mice; $t_{10}$ = 14.35, ****p < 0.0001.

(i) A representative example of visualizing the 3D-linear SVM (LSVM) where the activities of $IL^{\rightarrow NAcSh}$ neurons decode the identity of novel mouse.

(j) A representative example of visualizing the 3D-LSVM where the activities of $IL^{\rightarrow NAcSh}$ neurons decode the position of novel mouse.

(k) Both identity and position information of novel social targets were well classified using the 3D-LSVM model compared to those from null models. Identity, Two-tailed paired t-test: n = 11 mice; $t_{10}$ = 23.17, ****p < 0.0001. Position, Two-tailed paired t-test: n = 11 mice; $t_{10}$ = 28.10, ****p < 0.0001.

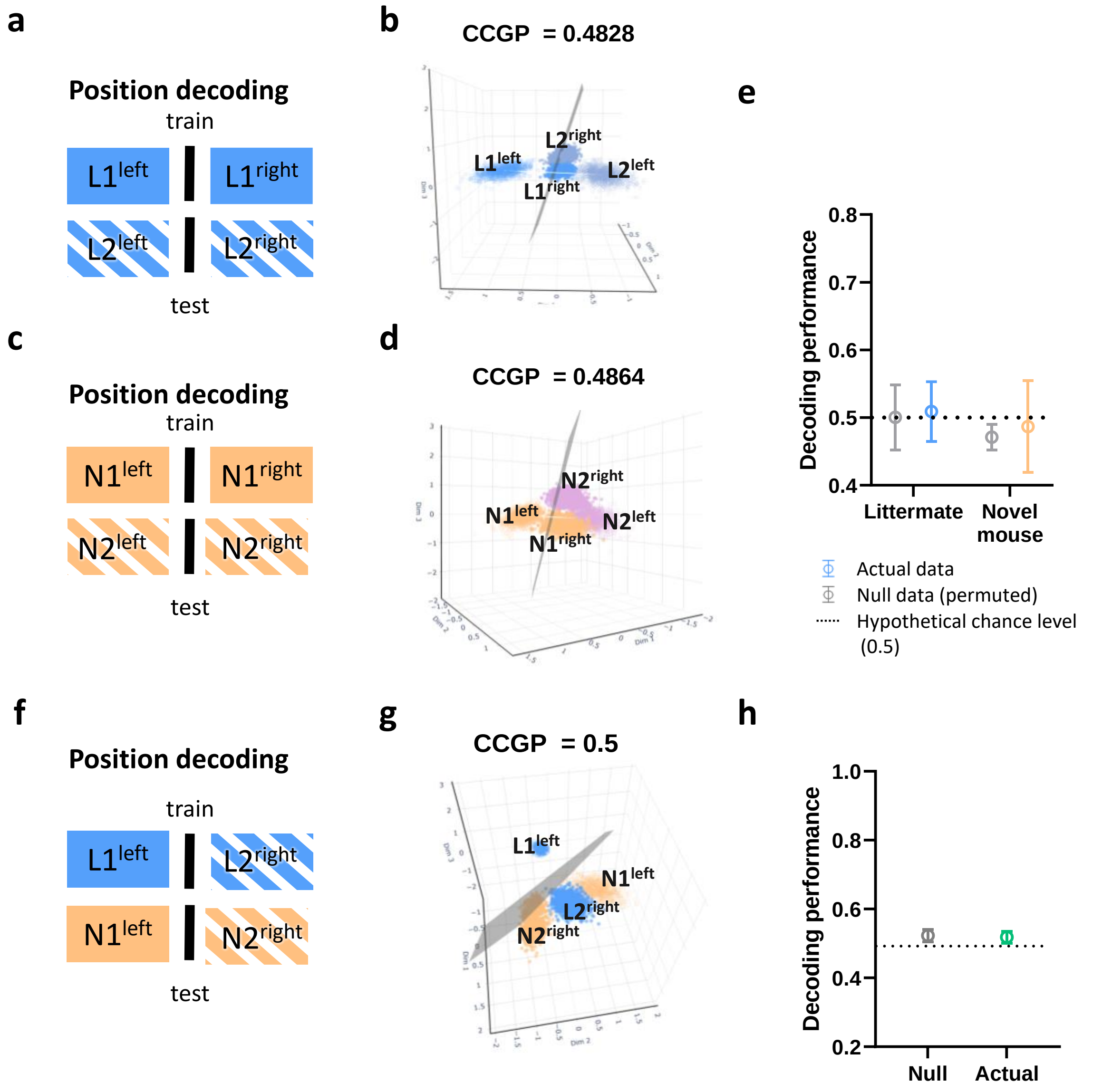


### Supplementary Figure 4.

**Supplementary Figure 4. $\text{IL}^{\rightarrow\text{NAcSh}}$ neural activity does not encode generalized spatial information of conspecifics**

(a) Schematic diagram for decoding position using CCGP classifier from the consecutive littermate recognition task.

(b) A representative image showing $\text{IL}^{\rightarrow\text{NAcSh}}$ neural activity does not support generalized decoding of littermate position.

(c) Schematic diagram for decoding position using CCGP analysis from the consecutive novel mouse interaction task.

(d) A representative image showing $IL^{\rightarrow NAcSh}$ neural activity does not support generalized decoding of novel mice position.

(e) Interaction-dependent $IL^{\rightarrow NAcSh}$ neural activity exhibited low CCGP performance scores for decoding positions comparable to those from the null models. Two-tailed paired t-test: n = 11 mice; Littermate $t_{10} = 0.3620$, $p = 0.7249$; Novel mouse $t_{10} = 0.2324$, $p = 0.8209$.

(f) Schematic diagram for decoding position using CCGP analysis from the littermates and novel mouse interaction task.

(g) A representative image showing $IL^{\rightarrow NAcSh}$ neural activity from littermates and novel mouse interaction for position CCGP analysis.

(h) Position CCGP analysis by comparing interaction-dependent $IL^{\rightarrow NAcSh}$ neural activity from littermate and novel mouse interactions exhibited a low decoding performance comparable to that from null data. Two-tailed paired t-test: n = 9 mice; $t_8 = 0.3225$, $p = 0.7553$.

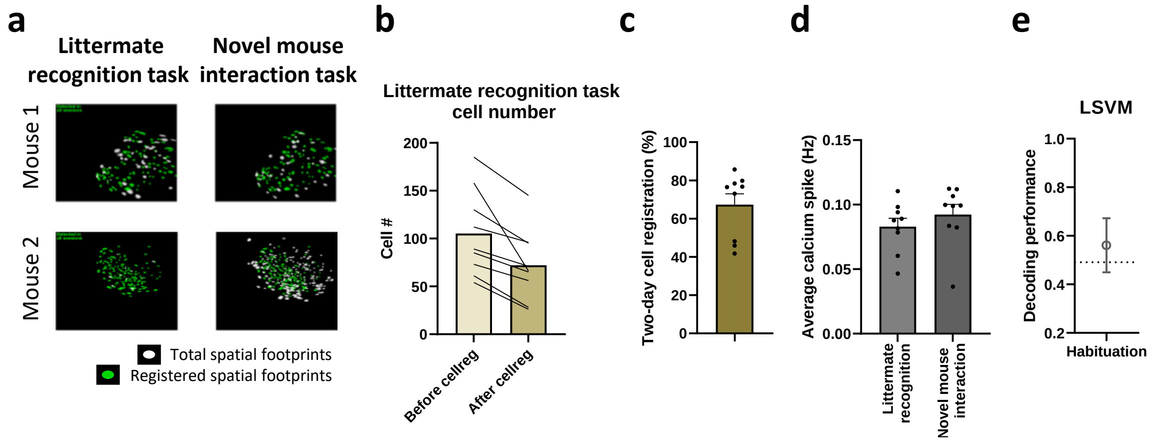


#### Supplementary Figure 5.

**Supplementary Figure 5. Registration of IL$^{\rightarrow NAcSh}$ neurons imaged in two different behavior tasks**

(a) Representative images from two subject mice showing longitudinal cell registrations from ROIs across multiple tasks. Among the total spatial footprints (white), footprints marked with green color represent registered cells..

(b) Longitudinal cell registration resulted in identification of smaller but significantly reduced cell populations within littermate recognition task. Two-tailed paired t-test: n = 9 mice; $t_8$ = 4.203, **p = 0.0030.

(c) 67.33% of neuronal populations were registered across multiple tasks.

(d) Average $Ca^{2+}$ transients were comparable during the littermate recognition task and the novel mouse interaction task. Two-tailed paired t-test: n = 9 mice; $t_8$ = 1.255, p = 0.2450.

(e) Calcium data obtained from the habituation session of each behavior paradigm was trained and tested with LSVM, revealing the decoding performance to chance level, ascertain the influence of social targets. One sample t-test: n = 9 mice; $t_8$ = 0.5515, p = 0.5964.

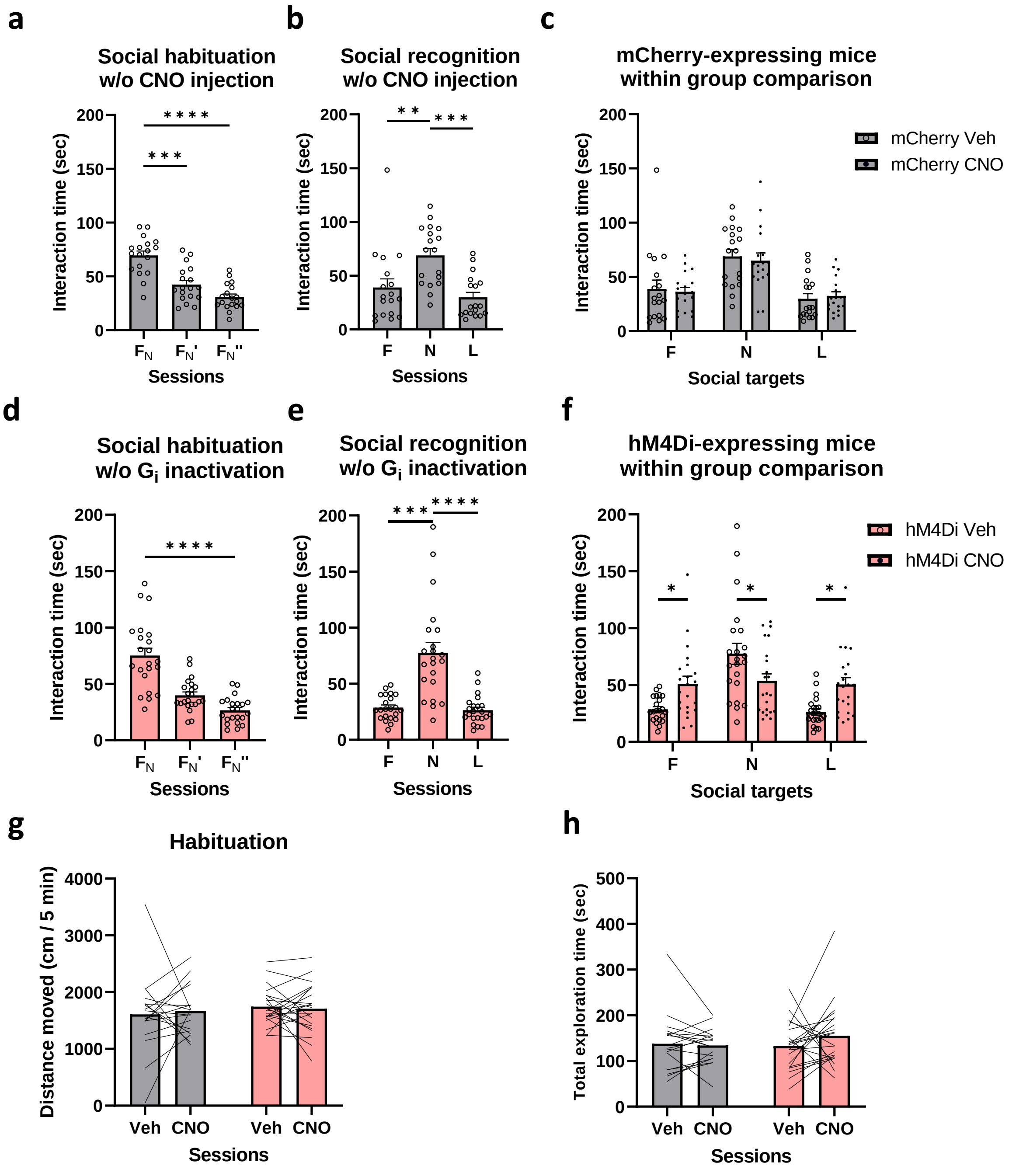


**Supplementary Figure 6.**

**Supplementary Figure 6. Control data for TRAP experiments**

(a) mCherry-expressing TRAPed mice without CNO injections showed a significant decreased interaction times during social familiarization session. One-way ANOVA with repeated measures: n = 18 mice; Effect of social target, $F_{1.828, 31.07}$ = 37.27, ****p < 0.0001. Tukey's multiple comparison test: $F_N$ versus $F_N$' $F_{17}$ = 7.823, ***p = 0.0001; $F_N$ versus $F_N$'' $F_{17}$ =

14.28, ****p <0.0001; $F_N$' versus $F_N$'' $F_{17}$ = 3.298, p = 0.0782.

(b) mCherry-expressing TRAPed mice without CNO injections showed a significant preference for novel conspecifics during the social recognition sesssion. One-way ANOVA with repeated measures: n = 18 mice; Effect of social target, $F_{1.670, 28.39}$ = 16.02, ****p < 0.0001. Tukey's multiple comparison test: F versus N $F_{17}$ = 5.259, **p = 0.0046; F versus L $F_{17}$ = 2.376, p = 0.2412; N versus L $F_{17}$ = 6.992, ***p = 0.0003.

(c) mCherry-expressing TRAPed mice showed comparable interaction times towards different social targets regardless of CNO injection. Two-way ANOVA with repeated measures: n = 18 mice; interaction between CNO effect and target effect, $F_{2, 51}$ = 0.2021, p = 0.8177. Sidak's multiple comparison test: mCherry Veh - CNO, F $t_{51}$ = 0.3503, p = 0.9798, N $t_{51}$ = 0.5006, p = 0.9446, L $t_{51}$ = 0.3422, p = 0.9811.

(d) hM4Di mice injected with saline showed significantly decreased interaction times during the familiarization session. Saline was injected after the familiarization session. One-way ANOVA with repeated measures: hM4Di, n = 18 mice; Effect of social target, $F_{1.471, 30.89}$ = 55.03, ****p < 0.0001. Tukey's multiple comparison test: $F_N$ versus $F_N$' $q_{21}$ = 9.661, ****p < 0.0001; $F_N$ versus $F_N$'' $q_{21}$ = 12.05, ****p < 0.0001; $F_N$' versus $F_N$'' $q_{21}$ = 6.092, ***p = 0.0009.

(e) Saline-injected hM4Di mice showed significantly longer interaction times towards novel conspecifics during the social recognition task. One-way ANOVA with repeated measures: hM4Di, n = 18 mice; effect of social target, $F_{1.179, 24.76}$ = 29.32, ****p < 0.0001. Tukey's multiple comparison test: F versus N $q_{21}$ = 7.372, ***p = 0.0001; F versus L $q_{21}$ = 1.013, p = 0.7566; N versus L $q_{21}$ = 8.481, ****p < 0.0001.

(f) Chemogenetic inactivation of TRAPed $IL^{\rightarrow NAcSh}$ neurons significantly altered interaction times with social targets. Two-way ANOVA with matching both factors: hM4Di, n = 22 mice; interaction between target effect and recognition-CNO effect, $F_{2, 63}$ = 11.56, ****p < 0.0001. Sidak's multiple comparison test: F, hM4Di Veh versus CNO $t_{63}$ = 2.797, *p = 0.0203; N, hM4Di Veh versus CNO $t_{63}$ = 2.982, *p = 0.0122; L, hM4Di Veh versus CNO $t_{63}$ = 3.010, *p = 0.0112.

(g) Both mCherry- and hM4Di-expressing TRAPed mice showed a comparable basal locomotor activity during familiarization session. Two-tailed paired t test: mCherry, n = 18 mice; $t_{17}$ = 0.3264, p = 0.7481, hM4Di n = 22 mice; $t_{21}$ = 0.3977, p = 0.6948.

(h) Both mCherry- and hM4Di-expressing TRAPed mice showed a comparable total exploration time during the social recognition task. Two-tailed paired t test: mCherry, n = 18 mice; $t_{17}$ = 0.3368, p = 0.7404, hM4Di n = 22 mice; $t_{21}$ = 1.233, p = 0.2313.

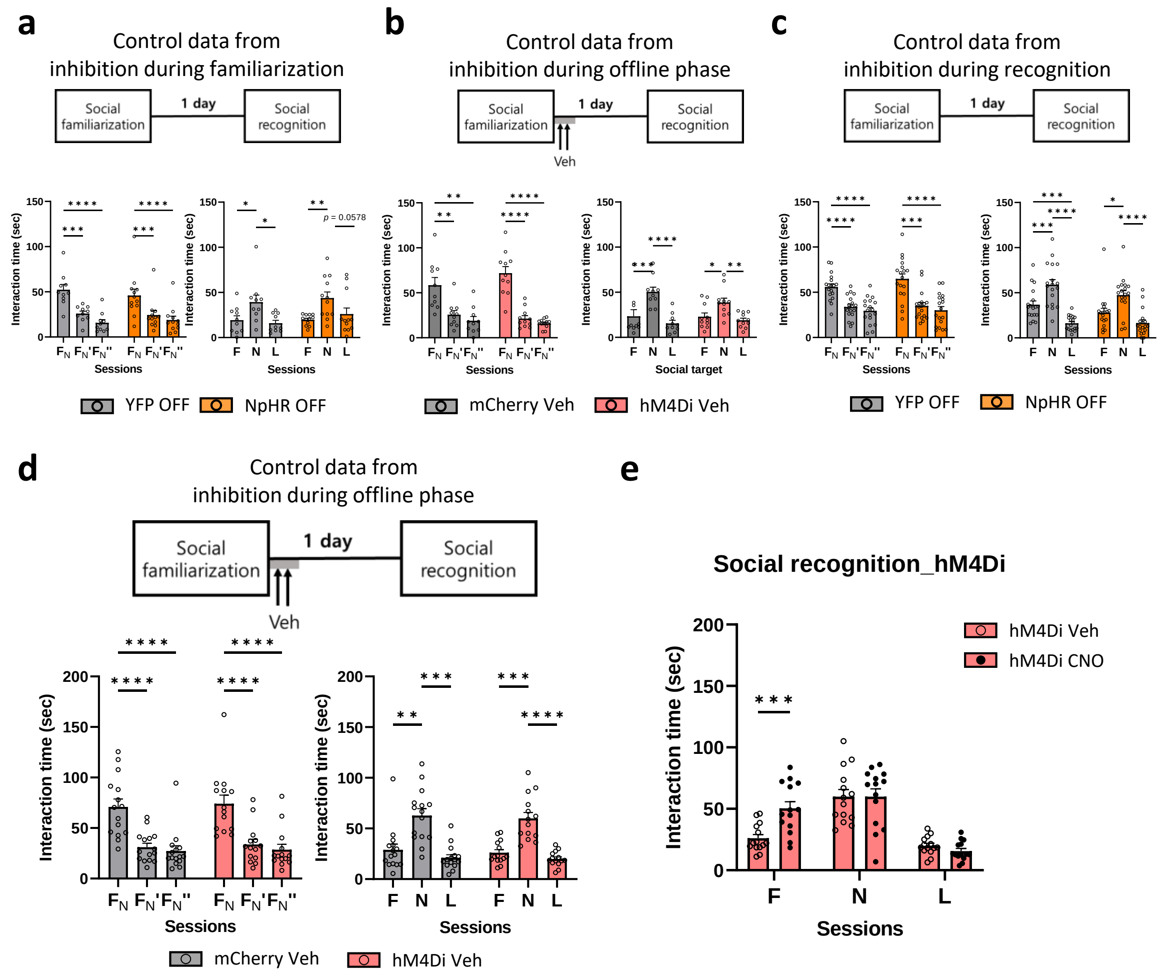


**Supplementary Figure 7.**

**Supplementary Figure 7. Control data for inactivating vCA1$^{\rightarrow IL}$ neurons**

(a) Both YFP and NpHR-expressing mice showed normal social recognition behaviors without laser delivery during social familiarization session. Left: social interactions during the social familiarization session. Two-way ANOVA with repeated measures: YFP, n = 10 mice; NpHR, n = 12 mice; interaction between session and NpHR expression effect, $F_{2, 40}$ = 0.6859, p = 0.5095. Tukey's multiple comparison test: vCA1 YFP, $F_N$ versus $F_N$' $q_{40}$ = 6.278, ***p = 0.0002; $F_N$ versus $F_N$'' $q_{40}$ = 8.710, ****p < 0.0001, $F_N$' versus $F_N$'' $q_{40}$ = 2.432, p = 0.2103; NpHR, $F_N$ versus $F_N$' $q_{40}$ = 5.659, ***p = 0.0008, $F_N$ versus $F_N$'' $q_{40}$ = 7.085, ****p <0.0001, $F_N$' versus $F_N$'' $q_{40}$ = 1.426, p = 0.5761. Right: social interactions during the social recognition session. Two-way ANOVA with repeated measures: YFP, n = 10 mice; NpHR, n = 12 mice; interaction between session and NpHR expression effect, $F_{2, 60} = 0.3785$, p = 0.6865. Tukey's multiple comparison test: YFP, F versus N $q_{60} = 3.404$, *p = 0.0495, F versus L $q_{60} = 0.5822$, p = 0.9110, N versus L $q_{60} = 3.987$, *p = 0.0177; NpHR, F versus N $q_{60} = 4.490$, **p = 0.0066; hM4Di, F versus L $q_{60} = 1.179$, p = 0.6836, N versus L $q_{60} = 3.311$, p = 0.0578.

(b) Both mCherry and hM4Di mice showed normal social recognition behaviors without CNO injections. Left: social interactions during the social familiarization session with saline injections. Two-way ANOVA with repeated measures: mCherry, n = 10 mice; hM4Di, n = 11 mice; interaction between session and hM4Di expression effect, $F_{2, 38} = 2.099$, p = 0.1366. Tukey's multiple comparison test: mCherry, $F_N$ versus $F_N$' $q_9 = 6.326$, **p = 0.0040, $F_N$ versus $F_N$" $q_9 = 5.912$, **p = 0.0060, $F_N$' versus $F_N$" $q_9 = 2.039$, p = 0.3615; hM4Di, $F_N$ versus $F_N$' $q_{10} = 10.28$, ****p < 0.0001, $F_N$ versus $F_N$" $q_{10} = 10.18$, ****p < 0.0001, $F_N$' versus $F_N$" $q_{10} = 2.594$, p = 0.2082. Right: social interactions during the social recognition session with saline injections. Two-way ANOVA: mCherry, n = 10 mice; hM4Di, n = 11 mice; interaction between session and hM4Di expression effect, $F_{2, 57} = 1.413$, p = 0.2518. Tukey's multiple comparison test: mCherry, F versus N $q_{57} = 5.657$, ***p = 0.0005, F versus L $q_{57} = 1.641$, p = 0.4815, N versus L $q_{57} = 7.298$, ****p <0.0001; hM4Di, F versus N $q_{57} = 3.456$, *p = 0.0458, F versus L $q_{57} = 0.8853$, p = 0.8065, N versus L $q_{57} = 4.341$, **p = 0.0091.

(c) Both YFP and NpHR mice showed normal social recognition behaviors without yellow laser delivery during social recognition session. Left: social recognition behaviors during the social familiarization. Two-way ANOVA with repeated measures: YFP, n = 18 mice; NpHR, n = 18 mice; interaction between session and NpHR expression effect, $F_{2, 68} = 1.103$, p = 0.3376. Tukey's multiple comparison test: YFP, $F_N$ versus $F_N$' $q_{17} = 8.553$, ****p < 0.0001, $F_N$ versus $F_N$" $q_{17} = 11.15$, ****p < 0.0001, $F_N$' versus $F_N$" $q_{17} = 1.996$, p = 0.3574; NpHR, $F_N$ versus $F_N$' $q_{17} = 6.759$, ***p = 0.0005, $F_N$ versus $F_N$" $q_{17} = 9.780$, ****p < 0.0001, $F_N$' versus $F_N$" $q_{17} = 1.488$, p = 0.5553. Right: social interactions during the social recognition session without laser delivery. Two-way ANOVA with repeated measures: YFP, n = 18 mice; NpHR, n = 18 mice; interaction between session and NpHR expression effect, $F_{2, 68} = 1.515$, p = 0.2271. Tukey's multiple comparison test: YFP, F versus N $q_{17} = 6.683$, ***p = 0.0005, F versus L $q_{17} = 7.203$, ***p = 0.0003, N versus L $q_{17} = 12.64$, ****p

<0.0001; NpHR, F versus N $q_{17} = 4.488$, *$p = 0.0146$, F versus L $q_{17} = 3.136$, $p = 0.0967$, N versus L $q_{17} = 8.673$, ****$p < 0.0001$.

(d) Both mCherry and hM4Di mice showed normal social recognition behaviors with vehicle infusions into the IL. Left: social recognition behaviors during the social familiarization. Two-way ANOVA with repeated measures: mCherry, n = 15 mice; hM4Di, n = 14 mice; interaction between session and hM4Di expression effect, $F_{2, 54} = 0.02629$, $p = 0.9741$. Tukey's multiple comparison test: mCherry, $F_N$ versus $F_N$' $q_{14} = 8.768$, ****$p < 0.0001$, $F_N$ versus $F_N$'' $q_{14} = 10.05$, ****$p < 0.0001$, $F_N$' versus $F_N$'' $q_{14} = 1.180$, $p = 0.6887$; hM4Di, $F_N$ versus $F_N$' $q_{13} = 9.048$, ****$p < 0.0001$, $F_N$ versus $F_N$'' $q_{13} = 10.84$, ****$p < 0.0001$, $F_N$' versus $F_N$'' $q_{13} = 2.500$, $p = 0.2185$. Right: social recognition behaviors during the social recognition session. Two-way ANOVA with repeated measures: mCherry, n = 15 mice; hM4Di, n = 14 mice; interaction between session and hM4Di expression effect, $F_{2, 57} = 0.02342$, $p = 0.9769$. Tukey's multiple comparison test: mCherry, F versus N $q_{14} = 5.978$, **$p = 0.0023$, F versus L $q_{14} = 1.623$, $p = 0.5019$, N versus L $q_{14} = 7.773$, ***$p = 0.0002$; hM4Di, F versus N $q_{13} = 7.223$, ***$p = 0.0005$, F versus L $q_{13} = 3.463$, $p = 0.0705$, N versus L $q_{13} = 9.263$, ****$p < 0.0001$.

(e) Chemogenetic inactivation of $vCA1^{\rightarrow IL}$ neurons by infusing CNO into the IL during offline phase impaired recognition of familiar conspecifics. Two-way ANOVA with matching both factors: hM4Di, n = 14 mice; interaction between target effect and offline - CNO effect, $F_{2,52} = 7.553$, **$p = 0.0013$. Sidak's multiple comparison test: F, hM4Di Veh versus CNO $t_{78} = 3.781$, ***$p = 0.0009$; N, hM4Di Veh versus CNO $t_{78} = 0.007777$, $p > 0.9999$; L, hM4Di Veh versus CNO $t_{78} = 0.6739$, $p = 0.8768$. F, familiar conspecific; N, novel conspecific; L, littermate.

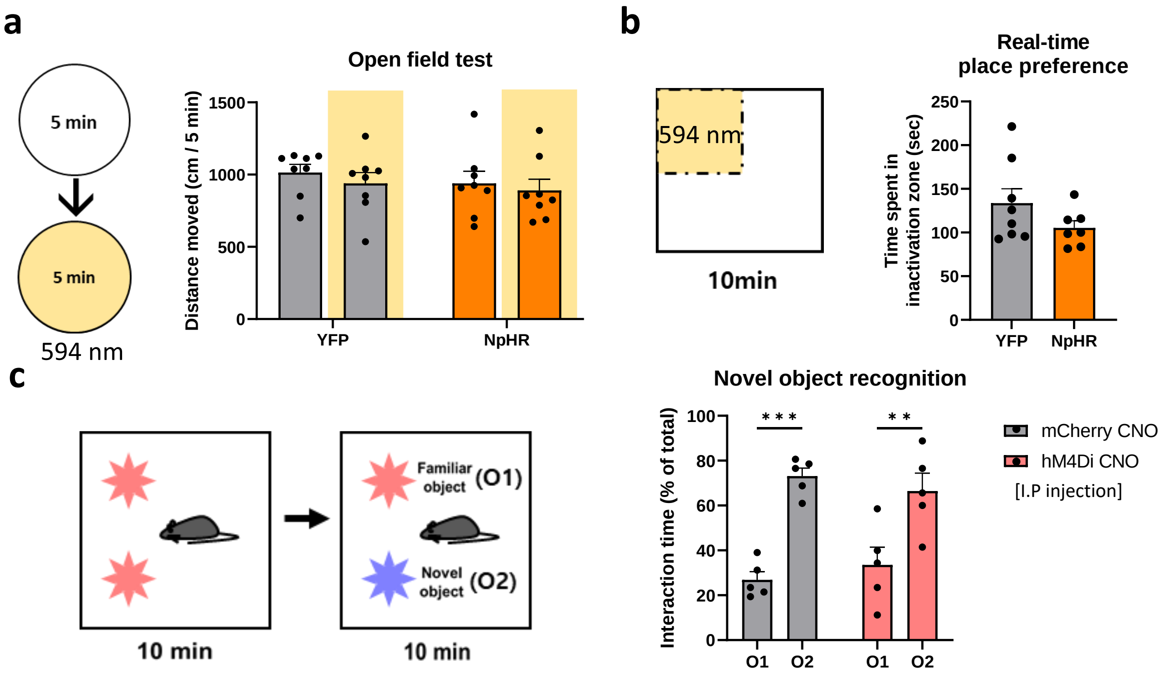


**Supplementary Figure 8.**

**Supplementary Figure 8. Inactivating vCA1$^{\rightarrow IL}$ neurons did not affect other behaviors**

(a) Left: schematic diagram of open field test (OFT) with NpHR inactivation. 594 nm laser with 6 mW intensity was given bilaterally to subject mice for 5 minutes. The order of light activation was counterbalanced. Right: inactivation of vCA1$^{\rightarrow IL}$ neurons did not affect mice basal locomotor activity. Two-way ANOVA with repeated measures: YFP, n = 8 mice; NpHR, n = 7 mice, $F_{1, 13} = 0.09891$, p = 0.7581. Sidak's multiple comparisons test: YFP, xlight – light $q_{13} = 0.9036$, p = 0.6189; NpHR, xlight – light $q_{13} = 0.4146$, p = 0.9009.

(b) Left: schematic diagram of real-time place preference (RTPP) with IL$^{\rightarrow NAcSh}$ neurons inactivation. Right: inactivation of IL$^{\rightarrow NAcSh}$ neurons did not affect place preference. Unpaired t-test: YFP, n = 8 mice; NpHR, n = 7 mice, $t_{13} = 1.458$, p = 0.1687.

(c) Left: schematic diagram of novel object recognition (NOR) with vCA1$^{\rightarrow IL}$ neurons inactivation. Right: both mCherry- and hM4Di-expressing mice showed a significant preference towards novel objects compared to familiar objects. Two-way ANOVA: mCherry n = 5 mice; hM4Di n = 5, interaction between CNO effect and hM4Di effect, $F_{1, 16} = 1.141$, p = 0.3013. Sidak's multiple comparison test: mCherry CNO, O1 versus O2 $t_{16} = 5.274$, ***p = 0.002, hM4Di CNO, O1 versus O2 $t_{16} = 3.763$, **p = 0.0034.

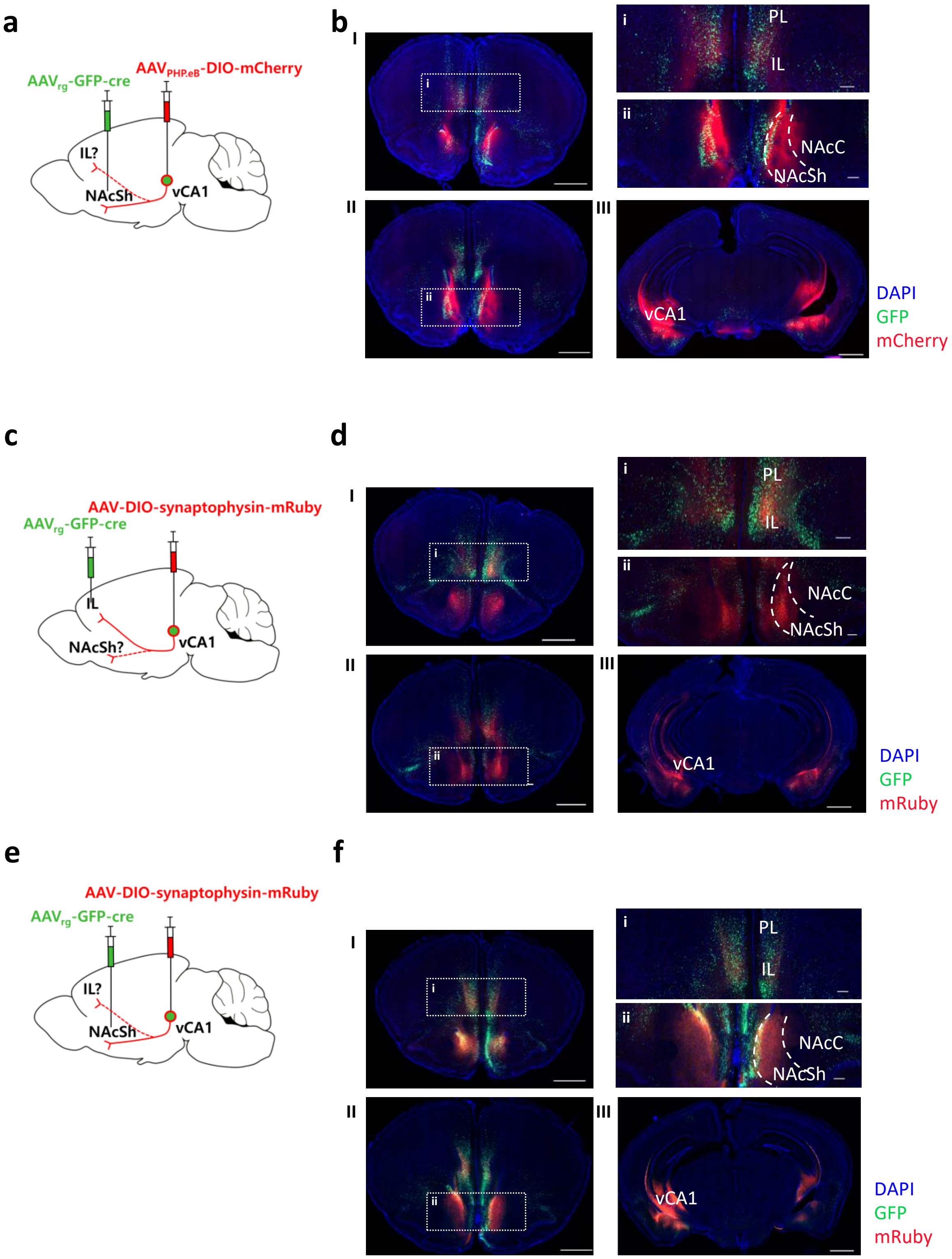


### Supplementary Figure 9.

**Supplementary figure 9. vCA1$^{\rightarrow IL}$ neurons exhibit collateral axonal terminals to NAcSh**

(a) Schematic diagram of labeling vCA1$^{\rightarrow NAcSh}$ neurons and their collateral projections.

(b) Axon projections were found in both IL (I and i) and NAcSh (II and ii) when vCA1$^{\rightarrow NAcSh}$ neurons were labeled (III). Scale bar: 1000 μm in whole brain coronal images and 200 μm in expanded images.

(c) Schematic diagram of labeling axon terminals of vCA1$^{\rightarrow IL}$ neurons and their collateral projections by expressing Cre-dependent synaptophysin-mRuby.

(d) Axon terminals were found in both IL (I and i) and NAcSh (II and ii) when vCA1$^{\rightarrow IL}$ neuronal terminal was labeled (III) with Cre-recombinase system. Scale bar: 1000 μm in whole brain coronal images and 200 μm in expanded images.

(e) Schematic diagram of labeling axon terminals of vCA1$^{\rightarrow NAcSh}$ neurons and their collateral projections by expressing Cre-dependent synaptophysin-mRuby.

(f) Axon terminals were observed in both IL (I and i) and NAcSh (II and ii) when vCA1$^{\rightarrow NAcSh}$ neuronal terminal was labeled (III) with Cre-recombinase system. Scale bar: 1000 μm in whole brain coronal images and 200 μm in expanded images.

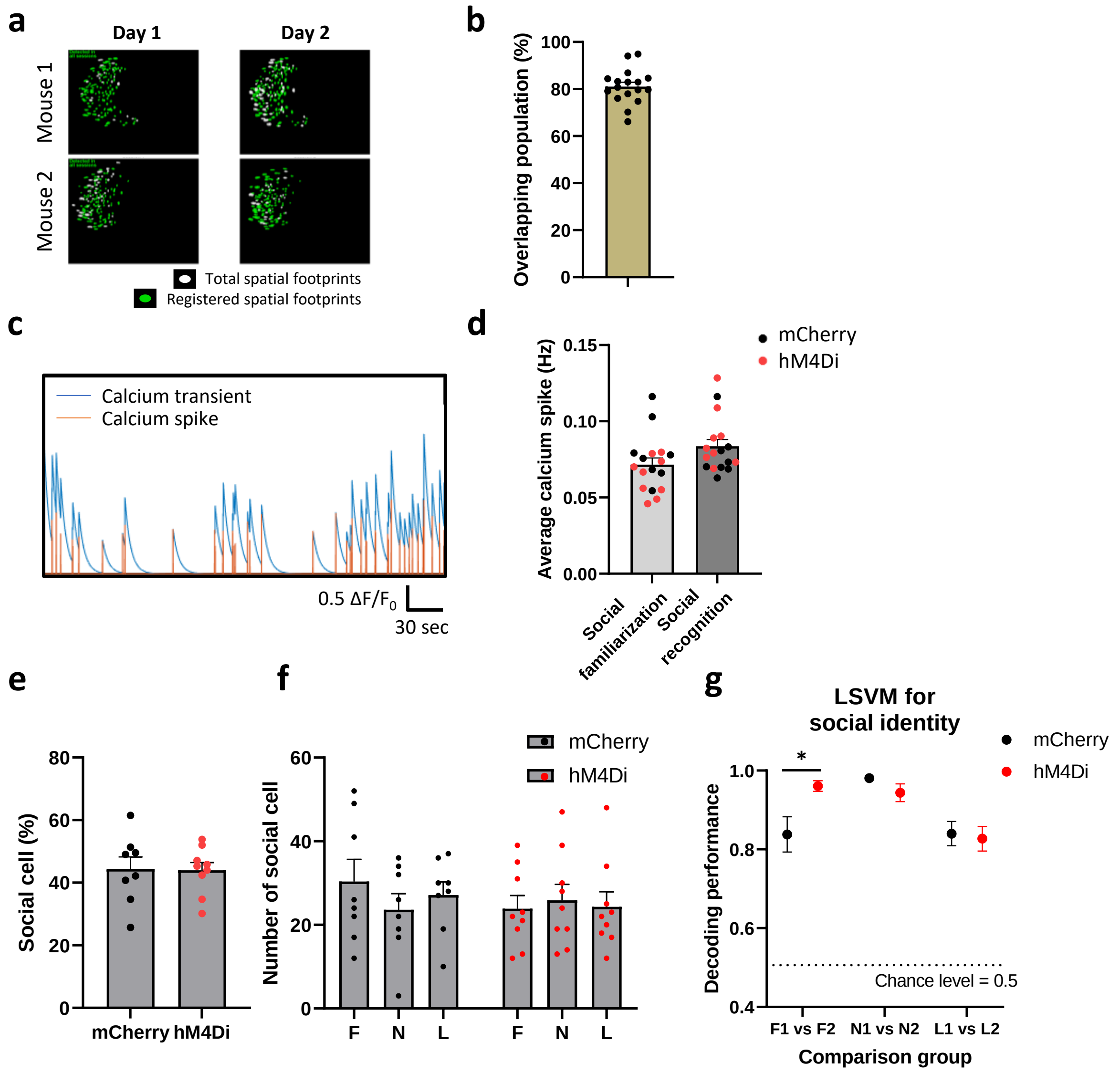


#### Supplementary Figure 10.

**Supplementary Figure 10. Imaging *in vivo* $Ca^{2+}$ activity of $IL^{\rightarrow NAcSh}$ neurons with vCA1 inactivation**

(a) Representative images from two subject mice showing longitudinal cell registrations from ROIs across multiple days.

(b) 81.07% of neuronal populations were registered across multiple days.

(c) Representative traces showing the $Ca^{2+}$ transient and deconvolved $Ca^{2+}$ spikes

(d) $Ca^{2+}$ transients during each habituation session was comparable between consecutive imaging days. Two-tailed paired t test: n = 14 mice; $t_{13} = 1.490$, p = 0.1600.

(e) mCherry- and hM4Di-expressing mice showed comparable size of social cell populations. Unpaired t-test, mCherry n = 8, hM4Di n = 9, $t_{15} = 0.08895$, p = 0.9303.

(f) Both mCherry- and hM4D-expressing mice exhibited comparable numbers of social cells for different social targets. Two-way ANOVA with repeated measures: mCherry n = 8, hM4Di n = 9, Interaction $F_{2, 30} = 2.245$, p = 0.1234, Tukey's multiple comparison test mCherry, F versus N, $q_{30} = 3.165$, p = 0.0810, F versus L, $q_{30} = 1.524$, p = 0.5351, N versus L, $q_{30} = 1.641$, p = 0.4855; hM4Di, F versus N, $q_{30} = 0.9946$, p = 0.7634, F versus L, $q_{30} = 0.2210$, p = 0.9866, N versus L, $q_{30} = 0.7736$, p = 0.8488.

(g) The LSVM classifier can successfully decode social identities of conspecifics with similar familiarity from neural activity during the social recognition session in both mCherry- and hM4Di-expressing mice. However, the performance for decoding two familiar conspecifics were significantly different between mCherry- and hM4Di-expressing mice. Unpaired t-test, F1 vs F2 mCherry versus hM4Di $t_{15} = 2.758$, *p = 0.0147.

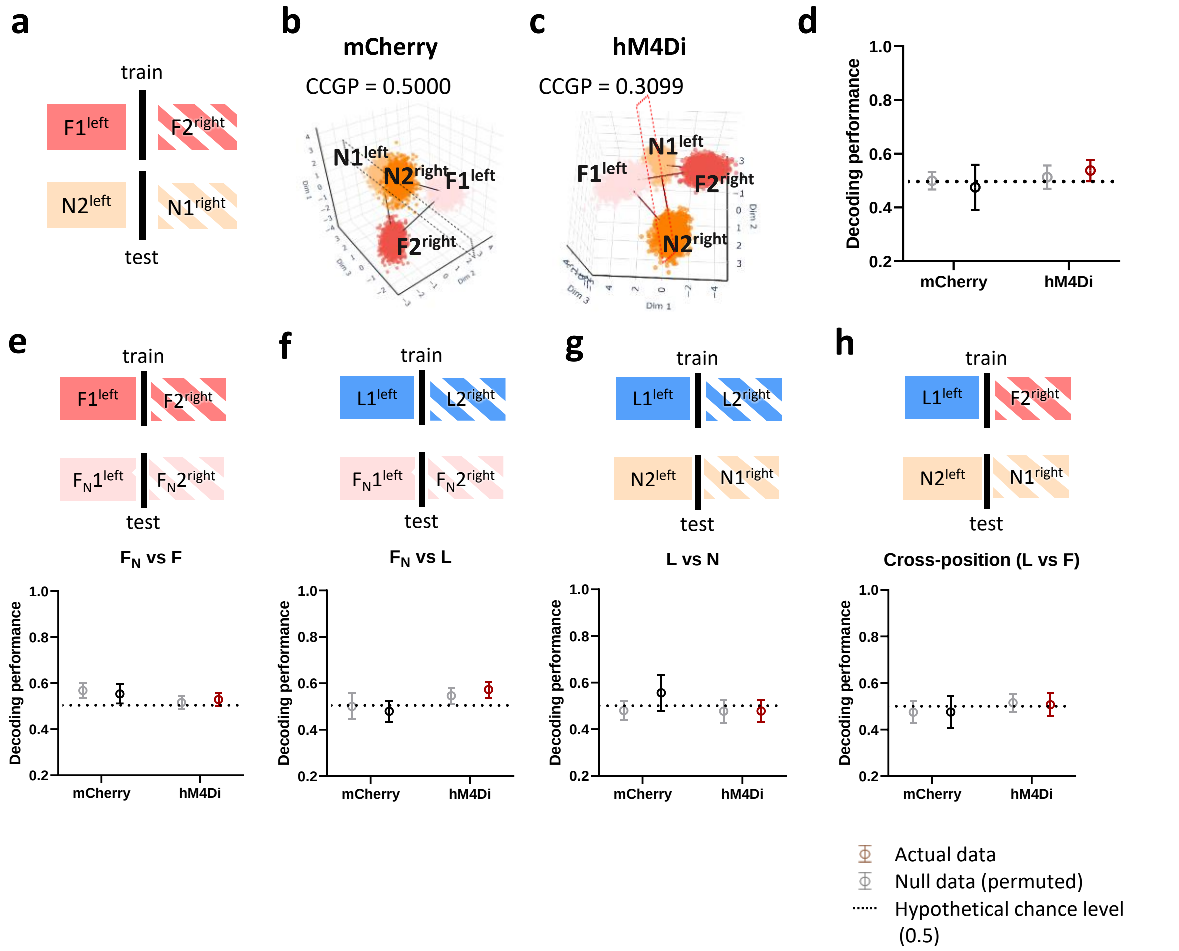


**Supplementary Figure 11.**

**Supplementary Figure 11. Inactivation of vCA1$^{\rightarrow IL}$ neurons did not affect position CCGP performance using IL$^{\rightarrow NAcSh}$ neural activity**

(a) Schematic diagram representing decoding for position CCGP between vCA1$^{\rightarrow IL}$ neuronal activities from familiar versus novel social targets.

(b) A representative geometry image of mCherry-expressing mice showing CCGP performance close to chance level for the position of conspecifics.

(c) A representative geometry image of hM4Di-expressing mice showing CCGP performance close to chance level for the position of conspecifics.

(d) Both mCherry and hM4Di-expressing mice showed chance levels of CCGP performance for position when comparing between familiar and novel social target. Two-way ANOVA with repeated measures: mCherry, n = 8 mice; hM4Di, n = 9 mice; interaction between hM4Di effect and null model effect, $F_{1,15}$ = 0.2319, p = 0.6371. Sidak's multiple comparison test: mCherry, Null versus Actual $t_{15} = 0.3351$, $p = 0.9335$; hM4Di, Null versus Actual $t_{15} = 0.3465$, $p = 0.9291$.

(e) Top: schematic diagram representing position CCGP decoding between littermates and novel conspecifics. Bottom: both mCherry- and hM4Di-expressing mice showed comparable CCGP performances to the null models. Two-way ANOVA with stacked matching: mCherry n = 8 mice, hM4Di n = 9 mice; interaction between hM4Di effect and null model effect, $F_{1, 15} = 0.4395$, $p = 0.5174$. Sidak's multiple comparison test: mCherry, Null versus Actual $t_8 = 0.9210$, $p = 0.6052$, hM4Di, Null versus Actual $t_9 = 0.01046$, $p > 0.9999$. $F_N$, unfamiliar conspecific

(f) Top: schematic diagram representing position CCGP decoding performances between littermates and familiar conspecifics. Bottom: both mCherry- and hM4Di-expressing mice showed comparable CCGP performances to the null models. Two-way ANOVA with stacked matching: mCherry n = 8 mice, hM4Di n = 9 mice; interaction between hM4Di effect and null model effect, $F_{1, 15} = 0.06943$, $p = 0.7958$. Sidak's multiple comparison test: mCherry, Null versus Actual $t_8 = 0.1131$, $p = 0.9922$, hM4Di, Null versus Actual $t_9 = 0.5040$, $p = 0.8568$.

(g) Top: schematic diagram representing position CCGP decoding performances between littermates and unfamiliarized conspecifics. Bottom: both mCherry- and hM4Di-expressing mice showed comparable CCGP performances to the null models. Two-way ANOVA with stacked matching: mCherry n = 8 mice, hM4Di n = 9 mice; interaction between hM4Di effect and null model effect, $F_{1, 15} = 0.4594$, $p = 0.5082$. Sidak's multiple comparison test: mCherry, Null versus Actual $t_8 = 0.4060$, $p = 0.9042$, hM4Di, Null versus Actual $t_9 = 0.5574$, $p = 0.8282$.

(h) Top: schematic diagram representing cross-position CCGP decoding performances between littermate and familiar conspecific with novel conspecifics. Bottom: both mCherry and hM4Di-expressing mice showed comparable CCGP performances to the null models. Two-way ANOVA with stacked matching: mCherry n = 8 mice, hM4Di n = 9 mice; interaction between hM4Di effect and null model effect, $F_{1, 15} = 0.01254$, $p = 0.9123$. Sidak's multiple comparison test: mCherry, Null versus Actual $t_{15} = 0.004227$, $p > 0.9999$, hM4Di, Null versus Actual $t_{15} = 0.1587$, $p = 0.9846$.
